# SPIN reveals genome-wide landscape of nuclear compartmentalization

**DOI:** 10.1101/2020.03.09.982967

**Authors:** Yuchuan Wang, Yang Zhang, Ruochi Zhang, Tom van Schaik, Liguo Zhang, Takayo Sasaki, Daniel Peric Hupkes, Yu Chen, David M. Gilbert, Bas van Steensel, Andrew S. Belmont, Jian Ma

## Abstract

Chromosomes segregate differentially relative to distinct subnuclear structures, but this genome-wide compartmentalization, pivotal for modulating genome function, remains poorly understood. New genomic mapping methods can reveal chromosome positioning relative to specific nuclear structures. However, computational methods that integrate their results to identify overall intranuclear chromosome positioning have not yet been developed. We report SPIN, a new method to identify genome-wide nuclear spatial localization patterns. As a proof-of-principle, we use SPIN to integrate nuclear compartment mapping (TSA-seq and DamID) and chromatin interaction data (Hi-C) from K562 cells to identify 10 spatial compartmentalization states genome-wide relative to nuclear speckles, lamina, and nucleoli. These SPIN states show novel patterns of genome spatial organization and their relation to genome function (transcription and replication timing). Comparisons of SPIN states with Hi-C sub-compartments and lamina-associated domains (LADs) from multiple cell types suggest constitutive compartmentalization patterns. By integrating different readouts of higher-order genome organization, SPIN provides critical insights into nuclear spatial and functional compartmentalization.

## MAIN TEXT

In human and other higher eukaryotic cells, interphase chromosomes are organized spatially within the cell nucleus (Kumaran et al., 2008; Bonev and Cavalli, 2016), such that their packaging and folding lead to dynamic interactions between genomic loci (Dekker et al., 2013). A key determinant of this intranuclear chromosome packaging are the interactions between chromosomes and heterogeneous constituents in the nucleus – in particular, nuclear compartments or nuclear bodies – including nuclear pore complexes, lamina, nucleoli, and nuclear speckles (Spector, 2006; Dundr and Misteli, 2010). Earlier work demonstrated the important connections between spatial localization of chromosome regions and gene expression regulation (Kumaran et al., 2008; Takizawa et al., 2008; Van Steensel and Belmont, 2017). Therefore, characterizing nuclear compartmentalization is crucial towards a comprehensive delineation of the roles of nuclear organization in different cellular conditions (Dekker et al., 2017). Unfortunately, our understanding of the genome-wide chromatin interaction with different nuclear compartments remains surprisingly limited.

The advent of whole-genome mapping methods for chromatin interactions such as Hi-C has shown that, at megabase resolution, chromosomes are spatially segregated into A/B compartments genome-wide (Lieberman-Aiden et al., 2009). A/B compartments exhibit distinct correlations to active euchromatic and inactive heterochromatic regions of the genome, respectively, although such strict, binary compartment separation is mostly partial (Kempfer and Pombo, 2019). Indeed, higher coverage Hi-C data generated from the human lymphoblastoid (GM12878) cells revealed that A/B compartments can be divided into five primary subcompartments, A1, A2, B1, B2, and B3, which harbor more refined associations with various functional features such as gene expression and histone modification (Rao et al., 2014). However, the observations of chromosome spatial association with nuclear compartments derived from Hi-C are limited and intrinsically indirect.

Several genome-wide mapping methods have enabled more direct examination between chromosome regions and specific nuclear compartments. In Guelen et al. (2008), DamID was utilized to measure contact frequencies between chromatin with nuclear lamina, revealing that∼35% of the human genome form lamina-associated domains (LADs). Recently, TSA-seq was developed to estimate cytological distance of chromatin toward nuclear speckles and nuclear lamina (Chen et al., 2018). Even though TSA-seq data show strong correlation with DamID, there are also clear differences. Most notably, the transitions of DamID scores are typically much more abrupt than TSA-seq maps that show gradual changes of signals over a chromatin trajectory (Chen et al., 2018), reflecting the differences in the methods, i.e., TSA-seq for molecular distance vs. DamID for contact frequency. In addition, Chen et al. (2018) (and later Xiong and Ma (2019)) showed that TSA-seq scores relative to speckles and lamina are correlated with Hi-C subcompartments although TSA-seq directly measures distance to the subnuclear structures. These results manifested the potential of an integrative framework that simultaneously analyzes different but complementary mapping data to offer a more complete view of nuclear compartmentalization.

Here, we develop a new computational method called SPIN (Spatial Position Inference of the Nuclear genome) to identify genome-wide chromosome localization patterns relative to multiple nuclear compartments. SPIN integrates TSA-seq, DamID, and Hi-C in a unified framework based on hidden Markov random field (HMRF). As a proof-of-principle, we apply SPIN to TSA-seq (for nuclear speckle and nuclear lamina), DamID (for nuclear lamina and nucleoli), and Hi-C data derived from K562 cells to identify genome-wide spatial localization states. The “SPIN states” reveal new and more detailed correlations with other features of genome structure and function, such as Hi-C subcompartments, topologically associating domains (TADs) (Dixon et al., 2012; Nora et al., 2012), histone modification, levels of transcriptional activity, and DNA replication timing. Comparisons with Hi-C subcompartments and LADs from multiple cell types suggest constitutive patterns of compartmentalization. We also identify possible molecular determinants and sequence-level features that modulate different compartmentalization. Taken together, SPIN is an effective integrative method that combines different genome-wide mapping approaches of nuclear genome organization to ascertain global patterns of spatial localization of the chromosomes, which provide critical insights into nuclear spatial and functional compartmentalization.

## Results

### Overview of the SPIN method

The overview of the SPIN method is illustrated in Fig. 1a. Our goal is to identify genome-wide spatial compartmentalization patterns of the chromosomes by integrating TSA-seq (Chen et al., 2018) and DamID (Guelen et al., 2008; Meuleman et al., 2013) data together with Hi-C. TSA-seq and DamID provide complementary information to measure distance and contact frequency between chromosome regions and subnuclear structures. The rationale of including Hi-C is that the pairwise genomic regions spatially interacting with each other more often than expected (from Hi-C) are more likely to share similar spatial compartmentalization patterns. We formulate this objective by using the hidden Markov random field (HMRF) (Zhang et al., 2001; Koller et al., 2009), in which nodes represent non-overlapping genomic bins (with a size of 25kb) and edges represent either significant Hi-C interactions or adjacent genomic bins (see Methods). We assume that each node is associated with an unobserved spatial localization state that SPIN aims to reveal (“SPIN state”). The observations on each node include signals from TSA-seq and DamID for defined nuclear compartments. Given the observed data on each genomic bin across the entire genome, the goal of SPIN is to solve the estimation problem by maximizing the likelihood of assigning spatial compartmentalization states. Thus the output of SPIN contains spatial localization state assignment, which is originally hidden, for each genomic bin throughout the genome.

**Figure 1:**
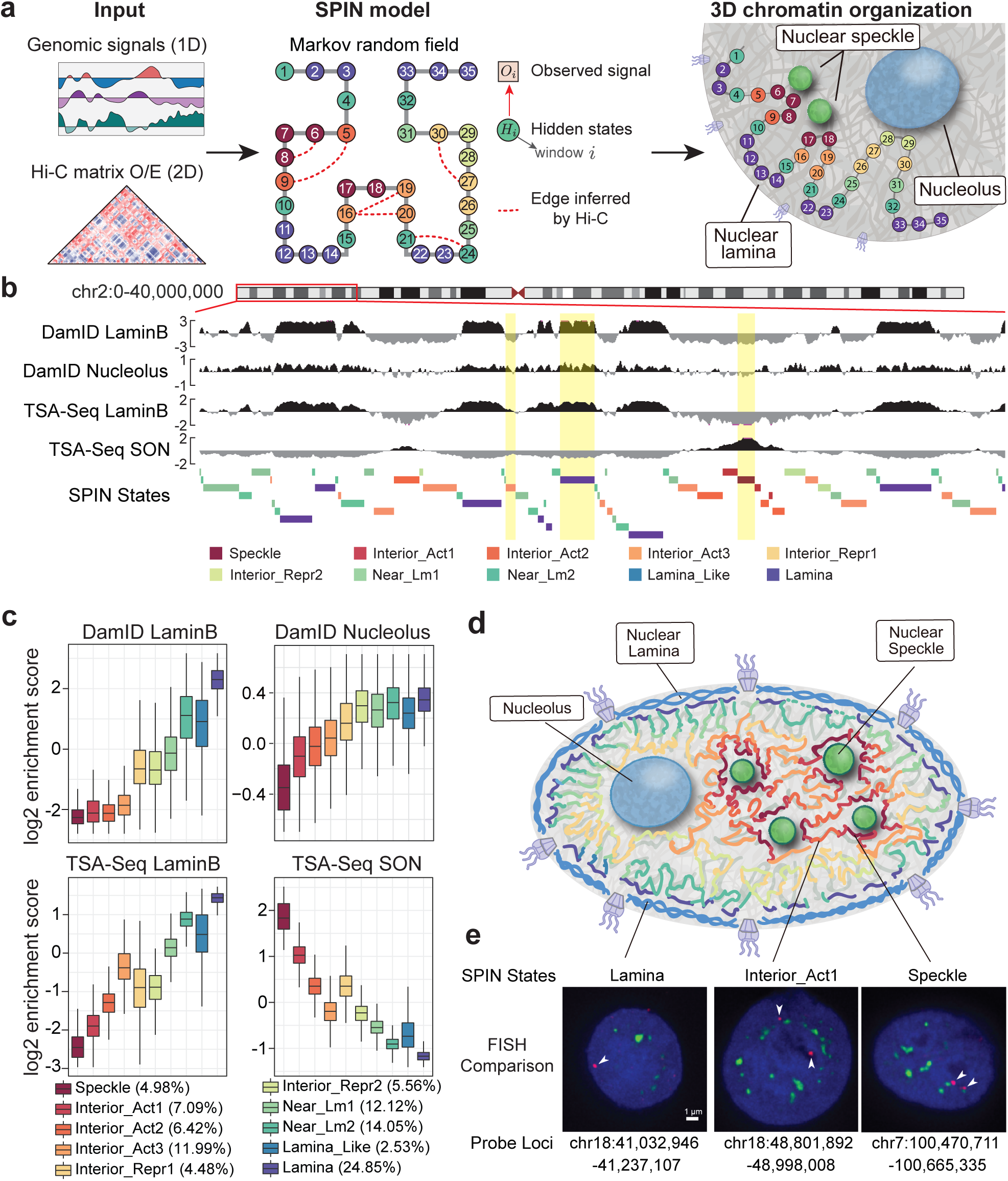
Overview of the SPIN method and the SPIN states in K562. **a.** Left panel: Input data for SPIN, including 1D genomic signals (TSA-seq and DamID), and Hi-C data. Middle panel: Graph representation of the hidden Markov random field. Genomic bins are represented as nodes and the hidden states of nodes are labeled by different colors. Right panel: Cartoon illustration of the SPIN states output for the nodes. **b.** Genome browser view of the input data (TSA-seq and DamID) and output (SPIN states). Three regions with different SPIN states (Interior _Act3, Lamina, and Speckle) are highlighted. These regions have distinct distributions of TSA-seq and DamID signals. **c.** Boxplots showing the distributions of the input TSA-Seq and DamID data on different SPIN states. The genome-wide percentage (size) for each SPIN state is also shown. **d.** Cartoon of the spatial positions of SPIN states relative to different nuclear compartments (nuclear lamina, nuclear speckles, nucleolus). **e.** Comparison between the identified SPIN states and DNA FISH. Three genomic regions with different SPIN states (Lamina chr18:41,032,946-41,237,107, Interior _Act1 chr18:48,801,892-48,998,008, and Speckle chr7:100,470,711-100,665,335) are shown. DNA probes are labeled in red (with white arrowhead) and nuclear speckles (SON protein) is labeled in green. Nuclear DNA staining (DAPI) is in blue.

SPIN is different from previous methods for chromatin domain segmentation based on a hidden Markov model (HMM) (Meuleman et al., 2013; Zheng et al., 2015; Marco et al., 2017) where chromatin interaction is not utilized. Although SPIN shares similarity in its goal with Segway-GBR (Libbrecht et al., 2015), the regularization in Segway-GBR uses significant Hi-C interactions as prior such that pairs of interacting genomic loci are encouraged to have the same label in genome annotation, which is not necessarily appropriate for refined spatial localization states of the chromosomes (see Supplementary Results for more detailed comparisons). In addition, the transition probabilities between different states learned in SPIN generalize such constraints so that chromatin regions in spatial proximity would have similar localization with more diverse patterns.

Note that in principle the input of SPIN on each genomic bin can also include functional genomic signals such as histone modifications, replication timing, and transcription levels. However, in this work we explicitly limit the input signals to those that directly measure the spatial position of chromatin (TSA-seq and DamID) and use functional genomic data to evaluate the functional correlations of different SPIN states genome-wide.

### SPIN identifies genome-wide patterns of nuclear compartmentalization

In this implementation of SPIN to infer genome-wide nuclear compartmentalization patterns, we used TSA-seq and DamID mapping data in K562 for nuclear speckles (TSA-Seq), lamina (DamID and TSA-Seq), and nucleoli (DamID). Nucleoli DamID data, generated using a Dam methylase fusion with a nucleolar targeting peptide repeat (4xAP3 (Scott et al., 2010)) is new; all remaining data are as published previously (Chen et al., 2018; Leemans et al., 2019). Details of the new nucleolar DamID mapping are described elsewhere (van Schaik et al. *manuscript in prep.*). Hi-C data for K562 are from Rao et al. (2014). Details of data processing for TSA-seq, DamID, and Hi-C are in the Supplementary Methods. We partition the genome (chromosome 1-22 and X) into consecutive non-overlapping 25kb bins, which constitute the graph structure for the HMRF model in SPIN. Edges are derived from Hi-C and the adjacent genomic bins (including those caused by large-scale structural variants in K562) (Methods and Supplementary Methods). Fig. 1b shows an example of the input signals from different measurements and the SPIN state output annotations.

We identified 10 SPIN states that represent major nuclear compartmentalization patterns in K562. These 10 SPIN states are: Speckle, Interior Active 1, 2, 3 (Interior _Act1, Interior _Act2, Interior _Act3), Interior Repressive 1, 2 (Interior _Repr1, Interior _Repr2), Near Lamina 1, 2 (Near _Lm1, Near _Lm2), Lamina Like, and Lamina. The genome-wide percentage of each state is shown in Fig. 1c. The name of these states are partially informed by comparing to various functional genomic data, especially for the Interior states (see later), even though the input for SPIN does not use any functional genomic data.

In Fig. 1c, we show that each SPIN state has distinct distributions of TSA-seq and DamID signals for the input data, reflecting the spatial position for compartmentalization. For example, the Speckle state has the highest SON TSA-seq signals and the lowest lamina/nucleolus signals as compared to other states. Notably, although we group multiple states into larger classes such as Interior Active, Interior Repressive, and Near Lamina, the refined states do show their distinct patterns. For example, the Interior _Repr2 state has similar DamID LaminB and TSA-seq LaminB as compared to Interior _Repr1, but its DamID nucleolus score is significantly higher while its SON TSA-seq score is significantly lower (*p*-value<2.2E-16). A recent report identified Nucleolus Associated Domains (NADs) in mouse embryonic fibroblasts and found that there are two types of NADs (Vertii et al., 2019): Type I NADs localize more frequently with both nucleoli and nuclear lamina and Type II NADs localize with nucleoli but do not overlap with lamina. We also observed such distinctions related to nucleoli from our SPIN states for spatial localization. The Interior _Repr2 state has similar enrichment of DamID nucleolous scores as compared to the Near _Lm1-2, Lamina Like, and Lamina states, but the Interior _Repr2 state has significantly lower enrichment with DamID LaminB and LaminB TSA-seq (Fig. 1c) (*p*-value<2.2E-16).

The identified SPIN states provide a comprehensive view of the spatial localization of the chromosomes in the nucleus relative to multiple subnuclear compartments, including nuclear speckles, nuclear lamina, and nucleoli (see the cartoon in Fig. 1d). We compared the SPIN states to DNA FISH (fluorescence *in situ* hybridization) imaging data. In Fig. 1e, we show three genomic regions (with detailed genomic coordinates) that correspond to different SPIN states with comparisons to DNA FISH data from Chen et al. (2018). The probe in the FISH image of the Speckle state has an average distance of 0.16*µ*m from nuclear speckle (SON protein). The probe in the FISH image of the Lamina state has an average distance of 0.98*µ*m from nuclear speckles and is located <0.5*µ*m from the nuclear periphery. The probe in the FISH image of the Interior Act1 state has an average distance of 0.47*µ*m from nuclear speckles. The comparison further suggests the reliability and advantage of having genome-wide SPIN states relative to multiple nuclear compartments.

### SPIN states provide spatial interpretation for Hi-C subcompartments

The five primary Hi-C subcompartments (A1, A2, B1, B2, and B3) defined from Rao et al. (2014), which exhibit strong associations with various genomic and epigenomic features, provide more detailed compartmentalization patterns from Hi-C data than the binary A/B compartment separation. However, the spatial localization context of Hi-C subcompartments has not been clearly revealed except that Chen et al. (2018) used the Hi-C subcompartments to identify the two transcriptional hot-zones based on TSA-seq scores by comparing to A1/A2 subcompartments (defined in GM12878 which has extremely high coverage Hi-C data). The recently developed algorithm SNIPER (Xiong and Ma, 2019) facilitates the identification of subcompartments in Hi-C data with low to moderate coverage and provides the Hi-C subcompartment annotations specifically in K562. Here we directly compare the 10 SPIN states with Hi-C subcompartments in K562.

Fig. 2a shows the overall comparison of Hi-C subcompartments in different SPIN states. Specifically, we found that the Speckle and Interior _Act1 states are strongly associated with A1 subcompartment (fold change enrichment 8.5 and 4.7, *p*-value<2.2E-16; Supplementary Methods). Interior _Act2 is strongly associated with A1, A2, and B1 subcompartments (fold change enrichment 3.8, 3.2, and 3.6, respectively, *p*-value<2.2E-16). Interior _Act3 is enriched with A2 subcompartment (fold change enrichment 3.1, *p*-value<2.2E-16). The Interior Repr1 and Interior Repr2 states have more B1 subcompartment (fold change enrichment 2.8, and 5.3, *p*-value <2.2E-16). We found that the Lamina _Like state is strongly enriched with B2 subcompartment (fold change enrichment 4.95, *p*-value<2.2E-16), while Lamina state is associated with both B2 and B3 subcompartments (fold change enrichment 3.16 and 5.58, *p*-value<2.2E-16). Together, different SPIN states have a strong correlation with different Hi-C subcompartments, supporting that Hi-C subcompartments reflect spatial positions relative to nuclear structures. However, the SPIN states offer a much more direct and refined interpretation of Hi-C subcompartments in terms of spatial compartmentalization. For example, although the Speckle, Interior _Act1, and Interior _Act2 states are all enriched with A1 subcompartment, they show distinguishable distributions regarding SON TSA-seq signals (Fig. 1c). This suggests that SPIN is able to further subdivide Hi-C subcompartment annotations into additional distinguishable spatial states of nuclear compartmentalization.

**Figure 2:**
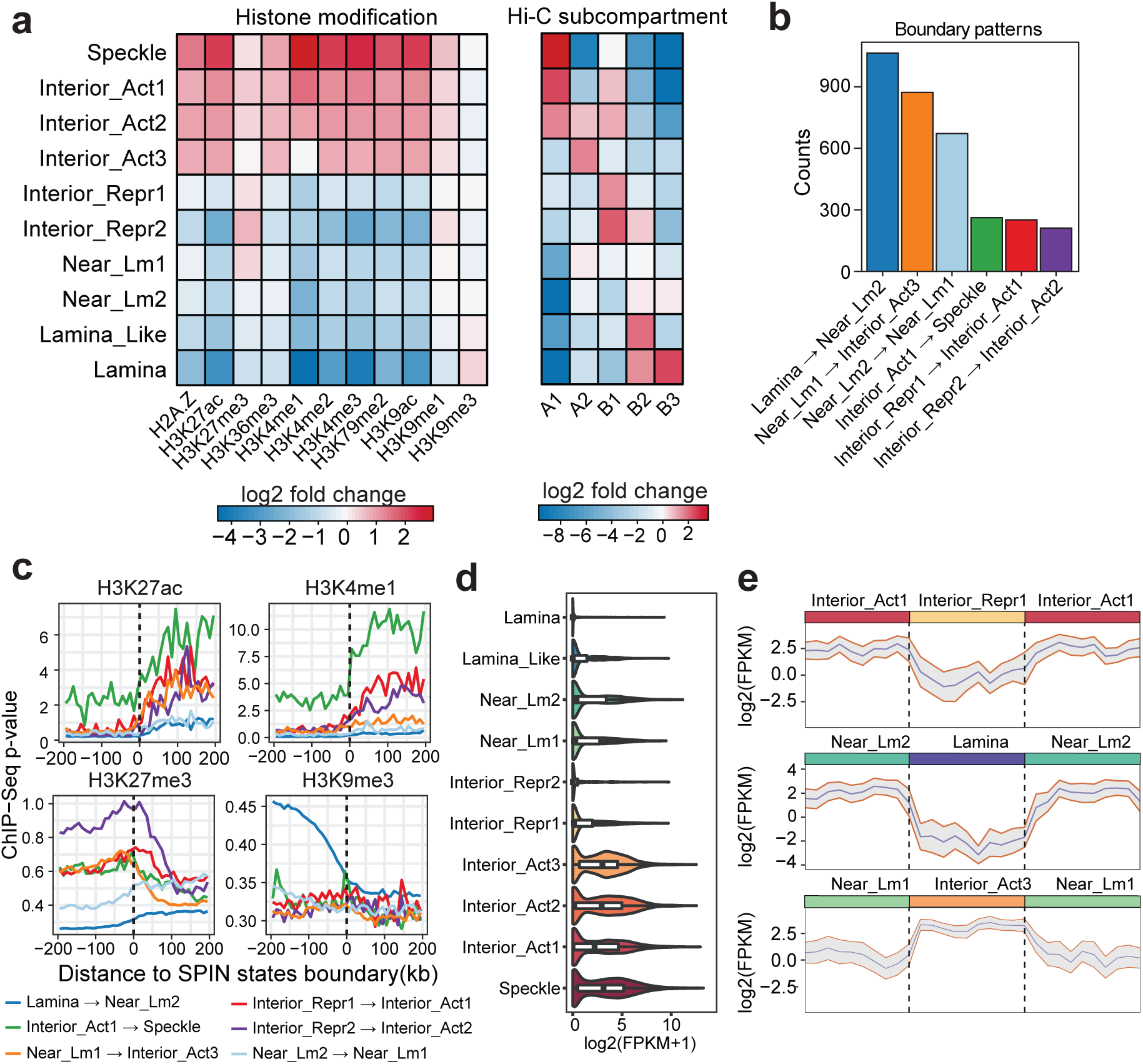
SPIN states correlate with various genomic and epigenomic features. **a.** Stratification of histone marks and Hi-C subcompartments as compared to SPIN states. Heatmap shows signal fold enrichmenh calculated as the ratio of observed signals over expected. K562 subcompartment annotations are produced by SNIPER (Xiong and Ma, 2019). **b.** Most frequently observed SPIN state boundary transition patterns. We count the number of different SPIN boundary patterns where symmetrical patterns are merged. **c.** Histone marks (H3K27ac, H3K4me1, H3K27me3, and H3K9me3) on SPIN state transition boundaries. **d.** Violin plot of transcript levels based on GRO-seq in different SPIN states. **e.** Transcript levels based on GRO-seq over three consecutive SPIN states. Three consecutive SPIN states are normalized by length (mean: blue line; standard deviation: gray).

### SPIN states stratify patterns of transcription activity and histone modification

Earlier studies have shown the correlation between the genome compartmentalization patterns and transcriptional activities (Lieberman-Aiden et al., 2009; Rao et al., 2014; Xiong and Ma, 2019). We sought to assess whether the SPIN states, which offer more detailed compartmentalization patterns, further stratify the transcriptional activity based on spatial locations of the chromatin. We first compared the SPIN states with 11 histone modification ChIP-seq datasets in K562 from the ENCODE project, including H3K79me2, H3K27ac, H3K9ac, H3K4me2, H3K4me3, H3K4me1, H3K9me3, H3K36me3, H2A.Z, H3K27me3, and H3K9me1. Overall, we found that the SPIN states have strong correlation with histone modifications (Fig. 2a). In addition, we also show that by adding Hi-C data into the SPIN model we can achieve state calling to better stratify histone modifications as compared to the baseline HMM-based model (Fig. S1). From the SPIN states, as chromatin localization changes from nuclear periphery to the interior (i.e., lamina to speckle axis), we observed a dramatic increase of ChIP-seq signal *p*-value of active histone marks (e.g., H3K27ac, H3K4me1, H3K4me3, H3K9ac) and, in general, a decrease in repressive marks (e.g., H3K9me3) (Fig. S2). Specifically, histone marks that are associated with transcriptional activation, including H3K4me1, H3K4me2, H3K4me3, H3K9ac, H3K27ac, and H3K79me2, show a dramatic increase of signal in the Speckle state (>5-fold increase on average, *p*-value<2.2E-16) (Fig. 2a). This result is consistent with previous studies that transcriptionally active chromatin regions are spatially localized preferentially near nuclear speckle and towards the interior (Chen et al., 2018; Van Steensel and Belmont, 2017). In contrast, heterochromatin mark H3K9me3 shows stronger presence in the Lamina state (*p*-value<2.2E-16), agreeing with earlier reports that LADs are often heterohchromatin with inactive genes (Meuleman et al., 2013; Van Steensel and Belmont, 2017; Zheng et al., 2015). In addition, the H3K27me3 mark, known to be associated with repressed transcription (Ferrari et al., 2014), is more abundant in Interior _Repr2 (*p*-value<2.2E-16) compared with the Near _Lm1 and Interior _Repr1 states (Fig. 2a). Importantly, we found that there is an increase of DamID nucleolus signals in Interior _Repr2 compared with the Interior _Repr1 and also the Interior Act states (Fig. 1c), suggesting a possible localization preference of Interior _Repr2 towards the nucleolus. This is in concordance with the recent report that H3K27me3 marks are enriched on Type II NADs (Vertii et al., 2019), which are found associated with nucleoli but not with LADs. Our results suggest that SPIN states reflect different associations with histone marks, and chromatin enriched for H3K27me3 and H3K9me3 have distinct spatial localization preferences.

To further demonstrate that the SPIN states clearly stratify functional genomic signals, we analyzed the patterns of histone modification signals across the transition boundaries between neighboring SPIN states. We selected the top six boundary types that are most frequently observed (Fig. 2b). Since SPIN states do not distinguish DNA strands, we therefore merged transition patterns in both directions on the genome. For example, Lamina to Near _Lm2 transition and Near _Lm2 to Lamina transition are considered as the same type of transition boundary for this analysis. For each transition type, we calculated the average histone modification signals at +/- 200 kb surrounding the transition boundaries. We found that many histone modifications show a clear, dramatic change across the SPIN state transition boundaries (Fig. 2c, Fig. S3). In particular, the active histone marks, such as H3K4me1 and H3K27ac, show a pronounced, >2-fold signal increase, when the chromatin trajectory is going from Interior _Act1 to Speckle. H3K9me1 signals exhibit a gradual rather than a sharp increase across the transition boundaries (Fig. S3). Additionally, we observed the opposite trend of signal enrichment across the boundaries for the repressive marks such as H3K27me3 and H3K9me3 (Fig. 2c). We further compared histone mark changes at the transition boundaries of SPIN states when the boundaries were defined by Hi-C subcompartments in K562 (Xiong and Ma, 2019). This reveals a sharper transition of histone marks at SPIN state boundaries as compared to Hi-C subcompartments (Fig. S3; especially H3K9me1, H3K9me3, H3K4me1, H3K4me2, H3K4me3), further suggesting that SPIN states offer a more accurate and refined definition of nuclear compartmentalization as compared to Hi-C subcompartments.

Next, we explored how transcription varies in different SPIN states. We compared SPIN states with GRO-seq data that measures through run-on transcription the density of engaged RNA Pol2 polymerases across protein coding genes in K562 (Fig. 2d). We found that genes in the Speckle and Interior Act states have high transcription levels, as expected, with average FPKM >40 for these run-on transcripts. The majority of the top 10% transcribed genes (over 90%) are from the Speckle or Interior Act states. In contrast, genes in the Lamina and Interior _Repr states are highly repressed. In addition, as shown in Fig. 2e for the consecutive SPIN states, transcription in Interior _Act vs. Interior Repr exhibit significant difference (t-test, *p*-value<2.2E-16), despite the fact that both states are likely to localize at relatively similar radial positions in the nucleus (based on TSA-seq). Also, genes in the Near _Lm and Lamina _Like states have higher transcription compared with the Lamina state (t-test, *p*-value = 5.689E-11). These analyses suggest that the SPIN states demarcate spatial patterns of chromosome regions in fine-scale separated into transcriptionally active and repressed regions.

### SPIN states are predictive of DNA replication timing

DNA replication timing (RT) is a vital genome function that is highly aligned with large-scale compartmentalization (Dileep et al., 2015). To further evaluate the functional connections of the SPIN states, we generated 7-fraction Repli-seq data for K562, where each fraction corresponds to the DNA replicated during 6 stages of S-phase as well as G2-phase representing the very latest DNA to replicate (Supplementary Methods). For each genomic bin (5kb), we calculated the percentage of DNA replicated in each fraction which was used to compute the fold change score of SPIN states on different replication fractions. We found that RT can be clearly stratified by the SPIN states (Fig. 3a). The Speckle and Interior Act states are found in early replicated regions (S1, S2, fold change score >1.5). The Interior _Repr1, Interior _Repr2, Near _Lm1 and Near _Lm2 states are replicated in the middle of S phase (S3, S4, fold change score >1.3). The Lamina Like and Lamina states are replicated late (S5, S6 and G2, fold change score >1.3). Overall, the SPIN states show a striking separation of the multi-fraction Repli-seq. In addition, using the definition of constitutive and developmentally-regulated replication domains (RDs) (Dileep et al., 2015), we found the SPIN states have distinct correlation with different patterns of constitutive and developmental RDs (Fig. 3b). Constitutive RDs can be further separated as constitutive early (CE) and constitutive late (CL) domains. We found that 85% of the genomic regions in Speckle state is CE and 55% Lamina state is CL. In contrast, other SPIN states contain a higher proportion of developmentally-regulated RDs. In Fig. 3c, we show that the SPIN states also correlate with evolutionary patterns of RT between human and non-human primates based on the annotations from Yang et al. (2018). Here the RT patterns are in five groups: early (all primate species have early RT), late (all primate species have late RT), weakly early (4 out of 5 species have early RT), weakly late (4 out of 5 species have late RT), and unconserved (the rest). Collectively, these analyses reveal a strong correlation between the detailed nuclear spatial compartmentalization identified by SPIN and the DNA RT program as well as its constitutive patterns in different cell types and across different species.

**Figure 3:**
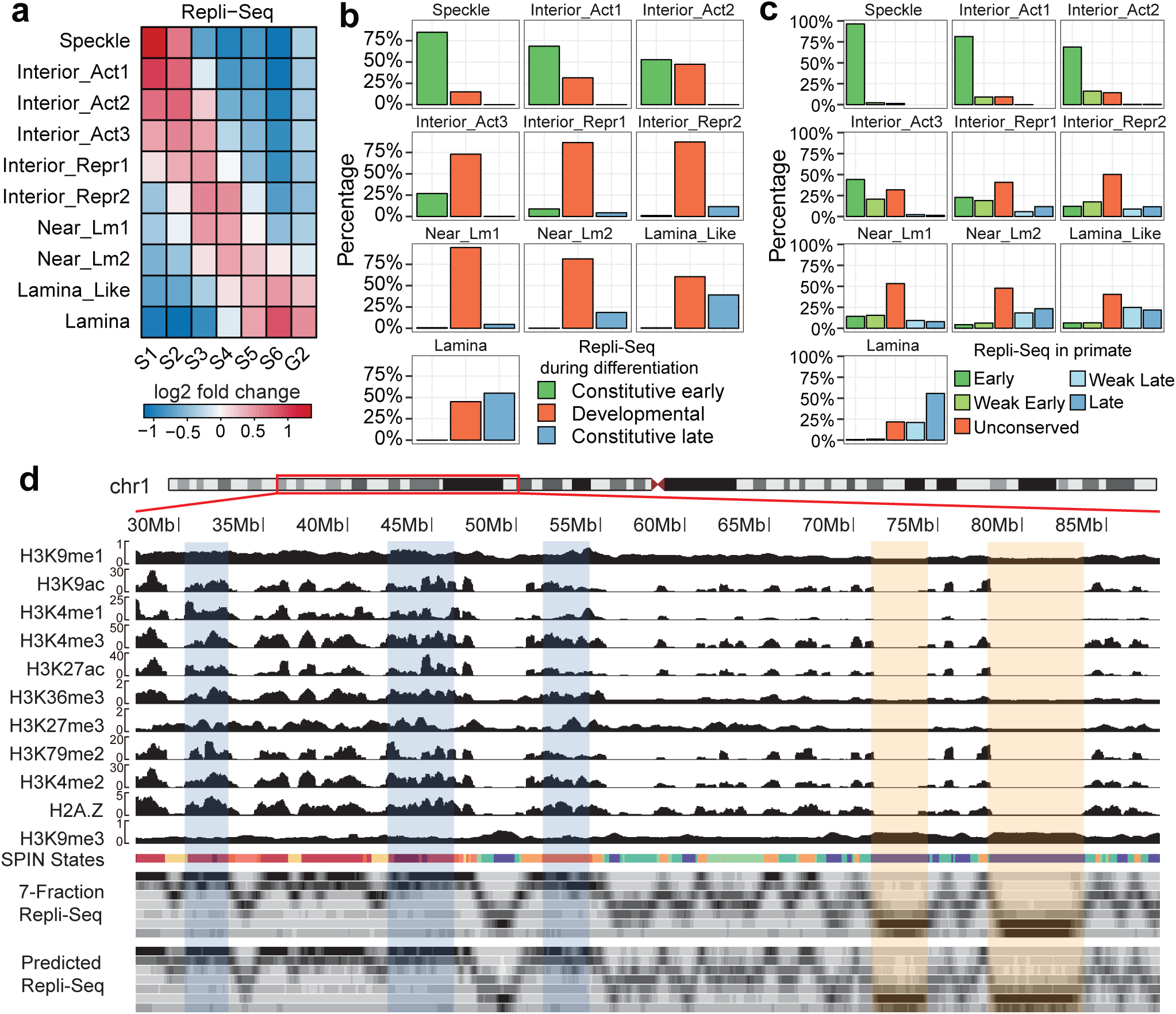
SPIN states are predictive of DNA replication timing. **a.** Stratification of multi-fraction Repli-Seq based on the SPIN states. **b.** Comparison with constitutive and developmentally-regulated replication timing (RT) domains. The barplot shows what percentage of each SPIN state belongs to constitutive early, constitutive late, and developmentally-regulated RT domains, respectively. **c.** Comparison with RT evolutionary patterns defined by Phylo-HMGP (Yang et al., 2018) between human and other non-human primate species. **d.** A Genome Browser view of the prediction of Repli-seq signals based on the SPIN states together with histone marks. The tracks are (from top to bottom): histone modifications, SPIN states, multi-fraction Repli-seq, and the predicted Repli-Seq signals. The three highlighted regions in light blue are examples of early replicating domains. The two highlighted regions in light orange are examples of late replicating domains.

Next, we sought to investigate the functional significance of SPIN states in terms of how important the SPIN states are, among other epigenomic features, in predicting RT. We built a predictive framework based on a random forest regression model to predict the multi-fraction Repli-seq signals along the genome by using the SPIN states together with various histone mark data (Fig. 3d) (see Methods). We specifically calculated the importance of each input feature based on how much each feature decreases the weighted impurity in a decision tree in the random forest. We found that the SPIN state is the most important feature, followed by H3K9me1, H3K9ac, H3K4me1, and H3K36me3, which are the top 5 most informative features (Fig. S4).

Together, the comparison with multi-fraction Repli-seq demonstrates that, by integrating different nuclear genome mapping data (TSA-seq, DamID, and Hi-C), the SPIN states delineate the detailed connections between nuclear compartmentalization and replication timing.

### SPIN states offer new perspectives for other nuclear organization units

We evaluated the significance of the SPIN states with respect to providing new insights for other nuclear genome features. We assessed the interplay between the SPIN states and the known 3D genome structural features, including LADs, TADs, and chromatin loops. We also asked whether the SPIN states are indicative of the constitutive patterns of nuclear compartmentalization across different cell types.

By combining TSA-seq and DamID, we identified several types of nuclear periphery states with different localization relative to nuclear lamina (Lamina, Lamina Like, and Near _Lm1-2, Fig. 1c). To further assess how each SPIN state corresponds to LADs across multiple cell types, we used DamID LaminB data in 6 human cell lines from the 4DN portal, including HCT116, K562. RPE-hTERT, HAP-1, HFFc6, and H1-hESC (see Table S1). Based on the assignment of LADs in 6 cell lines, we separated LADs into two categories: constitutive LADs (cLADs) and facultative LADs (fLADs). cLADs are defined as genomic regions characterized as LADs in at least 5 out of 6 cell lines. fLADs are defined as genomic regions characterized as LADs in at least 2 but fewer than 5 cell lines. In Fig. 4a, we show that there is a large difference between cLADs and fLADs in terms of the overlap with different SPIN states. 71% of cLADs are in the Lamina state in K562 as well as 22% in Near _Lm2, 3% in Lamina Like, and 3% in Near _Lm1 in K562. In contract, for fLADs, only 23% are in K562 Lamina state, but 39% are in Near _Lm2 and 23% are in Near _Lm1 in K562. This suggests that the SPIN states in one cell type (i.e., K562 in this study) can already separate fLADs and cLADs, as well as extending the concept of LADs into two separate categories. These results are consistent with the recently reported HiLands chromatin domains relative to the nuclear lamin based on both DamID LaminB and histone marks (in mES cells) (Zheng et al., 2015, 2018), where HiLands-B and HiLands-P are two distinct chromatin states that correspond to the facultative and constitutive LADs. For the two types of LADs, both the Lamina SPIN state and HiLands-P have higher DamID LaminB signals and higher H3K9me3 modification, while the Near _Lm SPIN state and HiLands-B have lower DamID LaminB signals and higher H3K27me3 enrichment.

**Figure 4:**
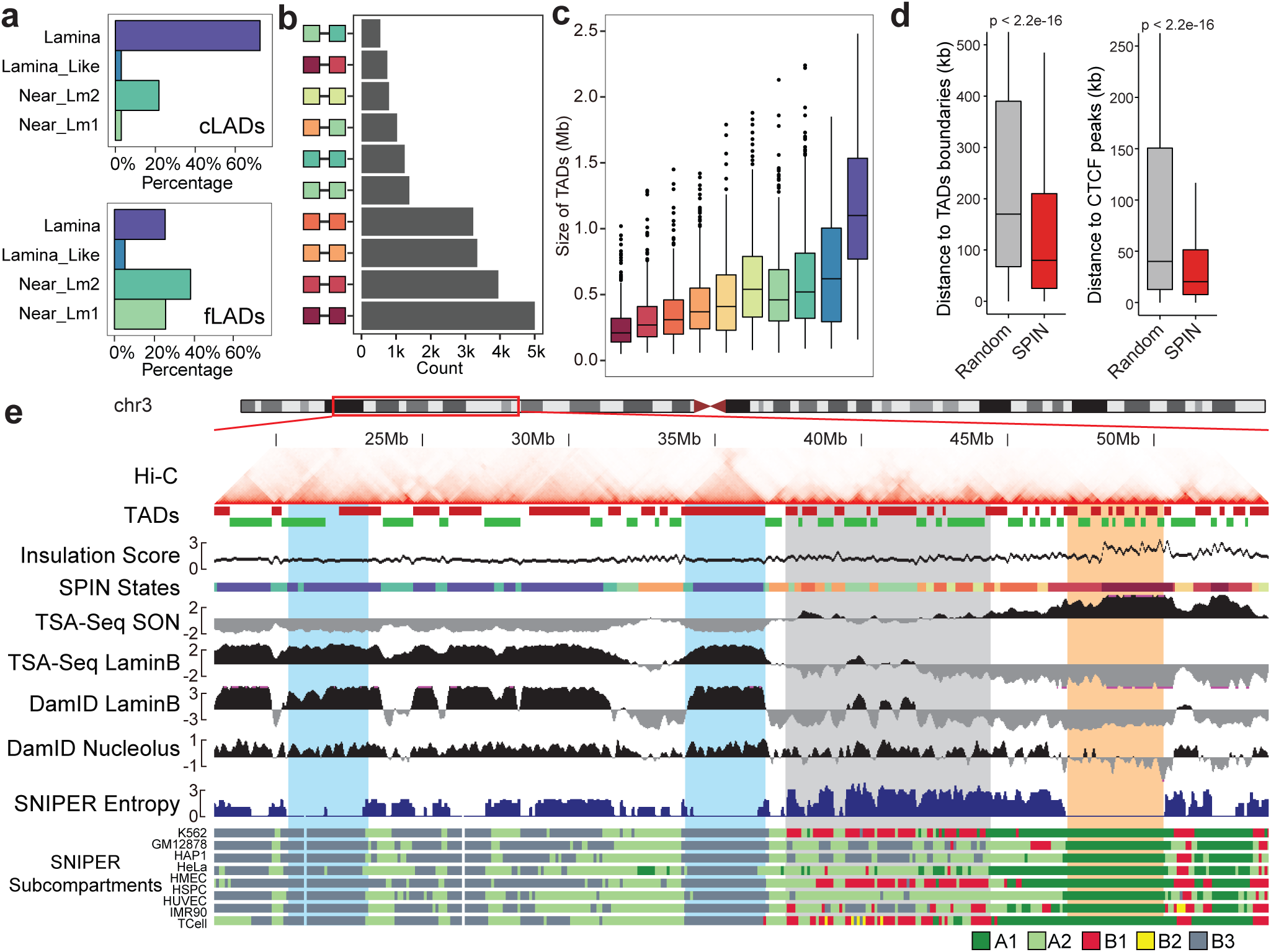
Comparison between the SPIN states and other nuclear organization units. **a.** Percentage of constitutive LADs (cLADs) and facultative LADs (fLADs) in SPIN states Lamina, Lamina Like, and Near _Lm1-2. cLADs and fLADs are called from DamID data from 6 different human cell lines. **b.** SPIN states of ChIA-PET significant interaction pairs (CTCF-mediated chromatin loops). **c.** Size of the TADs within each SPIN states. TADs are called by the Directional Index (DI) method. **d.** SPIN states boundaries are close to TADs boundaries and CTCF peaks. **e.** A Genome Browser snapshot of the SPIN states together with TADs and cross cell-type Hi-C subcompartments. K562 Hi-C contact map is shown at the top. TADs are called by the DI method. Hi-C subcompartments in multiple cell lines (Xiong and Ma, 2019) are shown at the bottom. Two highlighted regions in light blue are constitutive B3 subcompartments that correspond to the Lamina SPIN state. The light gray region has more dynamic subcompartment annotations in different cell types. The light orange region highlights a constitutive A1 subcompartment that corresponds to the Speckle SPIN state.

Next, we compared the SPIN states with ChIA-PET chromatin loops and TADs derived from Hi-C. For CTCF-mediated ChIA-PET chromatin loops, we discarded loops within 25kb range to focus on longer range interactions. We found that most loops are formed within the same SPIN states with more loops towards the interior states with higher transcriptional activity (Fig. 4b). We also observed similar results in Pol2-mediated ChIA-PET loops (Fig. S5). For TADs defined by the directionality index (DI) method (Dixon et al., 2012), we found that the TADs tend to stay within the same SPIN state. Specifically, 82.3% of TADs have only one SPIN state labeled. It is rare (0.4%) that one TAD spans more than two different SPIN states. Importantly, we observed the increase of TAD size when the SPIN state trajectory is changing from the nuclear interior to the periphery, with the average TAD size as 1.12Mb in the Lamina state and 0.19Mb in the Speckle state, respectively (Fig. 4c and Fig. 4e). The boundaries of the SPIN states are close to TAD boundaries and CTCF peaks than expected at random (Fig. 4d) (*p*-value <2.2E-16). We calculated the Hi-C insulation score (Lajoie et al., 2015) to represent TAD boundary strength. We found that the insulation score on the Speckle state is on average 2 times higher than the score in the Lamina state (Fig. 4e), indicating that there are stronger TAD/subTAD boundaries in the Speckle states (Supplementary Methods). In addition, we performed analysis for TAD-TAD level interactions and showed that TADs from the same SPIN state tend to form spatially separated cliques (Fig. S6 and Supplementary Results). Taken together, these results show that SPIN states stratify TADs and chromatin loops by providing spatial context.

We sought to analyze how conserved the spatial compartmentalization patterns are across different cell types. Here we use Hi-C subcompartments in multiple cell lines as an estimation of chromosome spatial localization in different cell types. As we have already shown, Hi-C subcompartments are highly correlated with SPIN states although the SPIN states provide more detailed and explicit compartmentalization views relative to subnuclear bodies. We compared the SPIN states in K562 with the SNIPER Hi-C subcompartments across 9 human cell types (Xiong and Ma, 2019), including K562, GM12878, HAP1, HeLa, HMEC, HSPC, HUVEC, IMR90, and TCell. We calculated the SNIPER Entropy score as the metric of conservation for Hi-C subcompartments across cell lines (Fig. S7). The Speckle state has the lowest SNIPER entropy score (0.1 on average), strongly suggesting that Hi-C subcompartments on Speckle (mostly A1) are largely conserved across cell lines (Fig. 4e). The Lamina Like and Lamina states (mostly B2 and B3 subcompartments) also have relatively low SNIPER entropy scores (0.75 on average). The most dynamic SPIN states across cell types are Interior _Repr2 and Near _Lm1. This comparison with cross-cell type SNIPER Hi-C subcompartments suggests that different SPIN states have distinct patterns across cell types with Speckle being the most conserved state.

## Discussion

In this work, we developed SPIN, a probabilistic graphical model that integrates TSA-seq, DamID, and Hi-C to provide a comprehensive view of genome-wide nuclear compartmentalization to nuclear speckles, nuclear lamina, and nucleolus. We identified 10 SPIN states in K562 based on TSA-seq and DamID data together with Hi-C. We showed that different SPIN states represent different spatial localization preferences within the nucleus. Further analysis indicates that SPIN states have strong correlation with and also better stratify other functional genomic features, such as histone modification, transcription activity, and DNA replication timing, suggesting that the detailed SPIN states have important relationships to genome function. The SPIN states also facilitate the identification of potential molecular determinants and sequence features that may play roles in modulating nuclear genome compartmentalization (see Supplementary Results, Figs. S8, S9, S10, and S11), paving the way for further experimental validation to pinpoint the mechanisms that give rise to specific compartmentalization. Our computational framework is flexible to incorporate more data for other nuclear structures (such as PML bodies, nuclear pores, and pericentromeric heterochromatin) when they become available to achieve even more complete characterization of nuclear compartmentalization. We therefore expect that SPIN has the potential to become an important method for revealing nuclear compartmentalization in different cellular conditions.

Nuclear compartmentalization analysis has primarily focused on A/B compartments based on Hi-C data. Although five major Hi-C subcompartments were revealed in (Rao et al., 2014) in GM12878 cells, such analysis has not been possible in other datasets with low to moderate coverage until recently (Xiong and Ma, 2019). Additionally, prior work on LADs and inter-LADs also made the binary distinction of chromatin domains associated with the nuclear lamina, where LADs and inter-LADs largely correspond to A/B compartment separations. Our SPIN states significantly advance our understanding of the detailed spatial localization patterns, much beyond binary separation of the chromatin domains in terms of nuclear compartmentalization. Importantly, we have demonstrated that SPIN states offer new spatial interpretation of Hi-C subcompartments, clarifying the compartmentalization patterns of specific subcompartments relative to nuclear speckles, the nuclear lamina and nucleolus. Besides, SPIN states reveal more refined compartmentalization patterns as compared to Hi-C subcompartments, supported by comparisons with other genomic and epigenomic features.

Our SPIN states can be further evaluated and compared using other analysis methods, e.g., polymer simulations (Nuebler et al., 2018), 3D genome structure population modeling (Hua et al., 2018), and integration between chromatin interactome and regulatory network (Tian et al., 2020). In addition, recently published new genome-wide mapping methods (Quinodoz et al., 2018; Beagrie et al., 2017; Zheng et al., 2019) and approaches for assessing chromatin interaction dynamics (Finn et al., 2019; Belaghzal et al., 2019) could be incorporated into our framework. In this work, we made additional attempt to reveal the patterns of consecutive SPIN states on the chromosomes to reveal potential chromatin fiber trajectories with distinct functions (see Supplementary Results, Fig. S12), which can be further validated by Oligopaints/OligoSTORM imaging (Beliveau et al., 2012; Nir et al., 2018; Bintu et al., 2018).

The molecular determinants that modulate the maintenance and movement of compartmentalization remain largely elusive. Earlier microscopy studies identified genes associated with chromatin targeting to specific nuclear structure (e.g., Hsp70 transgene (Khanna et al., 2014)). Very recent work from Falk et al. (2019) postulated the roles of molecular determinants for the global changes of chromosome compartmentalization, although the exact players have yet to be identified. Our SPIN states facilitate the identification of potentially important sequence features for specific compartmentalization, which provides promising tool to help elucidate the mechanisms that maintain and modulate compartmentalization. We made initial effort to identify sequence features enriched in different SPIN states (Supplementary Results, Figs. S8, S9, S10, S11). This can be further facilitated by SPIN compartmentalization states in other cell types to prioritize important sequence features. Such analysis can be validated by genome engineering experiments. Overall, SPIN represents an important step forward in developing integrative computational tools to offer new perspectives of spatial organization of the chromosomes in the nucleus and their interplay with various subnuclear structures.

## Methods

### Data acquisition, processing, and availability

#### TSA-seq and DamID data

K562 cells were obtained from the ATCC. For TSA-seq, the cells were cultured following the ENCODE project recommendations. For DamID, the cells were cultured according to 4DN guidelines following the ATCC recommendations. TSA-seq data generation of SON and lamin B was described and reported in Chen et al. (2018). DamID of lamin B1 data generation was described and reported in Leemans et al. (2019). For nucleolus DamID, a tandem repeat of 4 copies of the nucleolus targeting domain of AP3D1, linked with flexible GGSGG-linkers (4xAP3 in short (Scott et al., 2010)), was codon optimized for expression in human cells (IDT). NheI and SalI restriction sites were added on the flanks and used to replace the LMNB1 gene with the 4xAP3 repeat in the Dam expression vector. The 4xAP3 Dam vector was used to generate nucleolus contact data identical to LMNB1 DamID-seq (Leemans et al. (2019), van Schaik et al. *manuscript in prep.*). Sequencing reads from TSA-seq and DamID were first mapped to the human reference genome (hg38; chromosome 1-22 and X). For TSA-seq data, PCR duplicates were removed using Samtools (Li et al., 2009) (rmdup command with default parameters). Next, for TSA-seq data, we calculated the number of reads mapped in sliding 20kb windows with 1kb step size on the genome. The normalized TSA-seq enrichment score was calculated as the log2 ratio of read counts between TSA pull-down sample and the input normalized by sequencing depth (Chen et al., 2018). The TSA-seq score was further smoothed by using Hanning window of length 21 following the same smoothing approach used in Chen et al. (2018). For DamID data, scores were similarly calculated as the log2 ratio of mapped reads between Dam-target and Dam-only samples (Meuleman et al., 2013). The signal was then averaged on sliding 20kb windows with 1kb step size with additional smoothing by Hanning window of length 21. The smoothed TSA-seq and DamID signals were binned in 25kb resolution.

#### Hi-C data

We obtained the Hi-C data of K562 cells from (Rao et al., 2014). Both intra-chromosomal and inter-chromosomal interactions were processed at 25kb resolution and VC SQRT normalization was applied to the Hi-C contact matrices. Hi-C data extraction and normalization were performed using Juicer (Du-rand et al., 2016). For intra-chromosomal contact maps, we calculated the log2 ratio between observed (O) over expected (E) interactions (i.e., O/E) for each pair of interactions. The rationale is to consider genomic loci (not necessarily close on 1D distance) that share spatial localization with higher than expected genome-wide Hi-C interaction patterns to facilitate the identification of compartmentalization. For inter-chromosomal interactions, the expected number of interactions was set to be uniformly distributed between genomic loci on different chromosomes. For each chromosome, we fitted a Weibull distribution for Hi-C contacts and kept those interactions with *p*-value < 10^−5^ as significant interactions for subsequent steps as input to the SPIN method. For each inter-chromosomal interaction, we also used *p*-value < 10^−5^ as cutoff for significant interactions but for each pair (*i, j*) we required that all neighboring pairs between *i* ± 1 and *j* ± 1 should be also significant to increase the reliability of added edge.

#### Multi-fraction Repli-seq

Multi-fraction Repli-seq was performed using an extension protocol to the E/L Repli-seq (Marchal et al., 2018). Briefly, K562 interphase cells were labeled with BrdU and sorted into 7-fractions (S1, S2, S3, S4, S5, S6, and G2) (Supplementary Methods). Note that cells in G1 fraction were collected at the very early side of G1 peak and are sequenced without BrdU Immunoprecipitation (IP), therefore we used G1 fraction as a control to remove copy number and mappability bias. For each Repli-seq fraction, we mapped the sequenced reads to the reference genome hg38. Then we calculated read counts on 10kb sliding windows with 1kb step size. The total number of mapped reads were then normalized to 1 million read counts per fraction. The raw signals of each window were normalized by the signals from G1. Genomic bins with 0 mapped reads in G1 were considered as unmappable regions. For each 1kb bin, the replication timing signal was calculated as the percentage of the total signal over the seven fractions. Finally, the replication timing signals were also binned in 25kb resolution.

#### Other epigenomic data and annotations

We compared SPIN states with other epigenomic datasets, such as Hi-C subcompartments, TADs, LADs, ChIP-seq, and GRO-seq. For Hi-C subcompartments, we used the K562 Hi-C subcompartment annotation produced by SNIPER (Xiong and Ma, 2019). Hi-C subcompartments in additional cell types were also from Xiong and Ma (2019). K562 histone mark and transcription factor ChIP-seq data were obtained from the ENCODE project (Consortium et al., 2012). Datasets with replicates were merged. For data sets with no processed *p*-value available, we used MACS2 (Zhang et al., 2008) to calculate ChIP-seq *p*-values (for narrow peak call, command bdgpeakcall is used). We downloaded CTCF and POLR2A ChIA-PET in K562 from the ENCODE project. ChIA-PET reads were processed using ChIA-PET Tool (Li et al., 2010) with default parameters. As for TADs, we used the DI method (Dixon et al., 2012) to call TADs based on 10kb resolution Hi-C. In addition, we used CaTCH (Zhan et al., 2017) to call hierarchical TADs. To identify LADs, we used a hidden Markov model to identify LADs and inter-LADs from K562 DamID LaminB data (Guelen et al., 2008). LADs annotations in additional cell types were collected from 4DN data portal. See Supplementary Methods for additional data analysis details by comparing to SPIN states.

All datasets used in this work are listed in Table S1.

### Algorithm description of the SPIN method

#### Overall design of the model

SPIN (Spatial Position Inference of the Nuclear genome) is developed based on a type of probabilistic graphical model called hidden Markov random field (HMRF) (Zhang et al., 2001; Koller et al., 2009), with the goal to identify genome-wide patterns of nuclear compartmentalization by integrating TSA-seq, DamID, and Hi-C (see Fig. 1a for the model overview). HMRF can be represented as an undirected graph *G* = (*V, E*), where each node *i* ∈ *V* represents a non-overlapping 25kb genomic region and *E* represents the set of edges. For each node *i*, the observation *O*_*i*_ ∈ ℝ^*d*^ is a vector of 1D TSA-seq and DamID signals on the genomic bin. Specifically, in this study these observations are TSA-seq SON for nuclear speckle, TSA-seq LaminB for nuclear lamina, DamID LaminB for nuclear lamina, and DamID 4xAP3 nucleoli. The edges (*i, j*) ∈ *E* in the graph *G* represent: (1) Significant Hi-C interactions between two genomic loci (see Hi-C data processing); (2) Adjacent nodes on the chromosome; (3) Since K562 is a cancer cell lines, we specifically consider adjacencies introduced by large structural variations (see Supplementary Methods).

Each node *i* has a hidden state *H*_*i*_, which represents the spatial localization of genomic bin *i* relative to multiple nuclear compartments. *H*_*i*_ is only dependent upon *O*_*i*_ and the neighbors of *i*, i.e., *N* (*i*) = {*j*|*j* ∈ *V*, (*i, j*) ∈ *E*}. Given the number of states *k* (see below on how we estimate *k*), our goal is to estimate the hidden states *H*_*i*_ for all nodes that maximize the following joint probability:

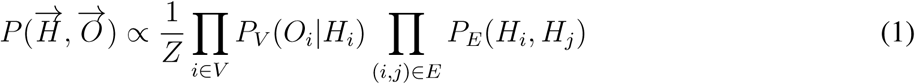

where 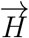 represents the hidden states of all nodes, *H*_*i*_ is the hidden state of node *i, P*_*V*_ (*O*_*i*_|*H*_*i*_) corresponds to the potential of node *i* that the observation is *O*_*i*_ given the hidden state *H*_*i*_. *P*_*E*_(*H*_*i*_, *H*_*j*_) corresponds to the edge potential between two nodes *i* and *j* with hidden states *H*_*i*_ and *H*_*j*_. *Z* is the constant used for normalization:

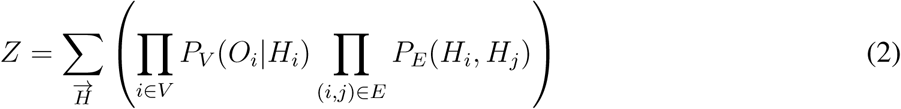

We assume that the observation of *O*_*i*_ given hidden state *H*_*i*_ = *h*_*a*_ follows a multivariate Gaussian distribution, i.e.,

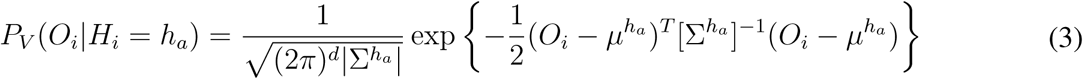

where *O*_*i*_ follows multivariate Gaussian distribution 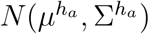 given state *H*_*i*_ = *h*_*a*_. The edge potential is defined by the transition probability between neighbor states *h*_*a*_ and *h*_*b*_.

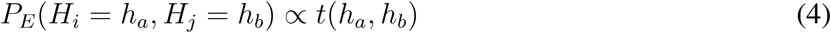

#### Initialization and model parameter estimation

To initialize *P*_*V*_ (*O*_*i*_|*H*_*i*_) for each node *i*, we estimate it based on a Gaussian mixture model with given number of states *k*. Here Gaussian mixture model assumes that the input data from TSA-seq and DamID for a given state are generated from a mixture of multivariate Gaussian distributions. We have:

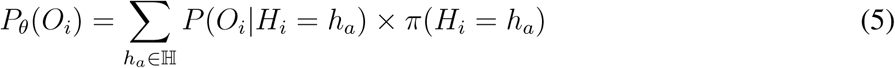

where *P*_*θ*_(*O*_*i*_) represents the mixture of *k* Gaussian distributions of different types of observed signals, *π*(*H*_*j*_ = *h*_*a*_) is the mixture proportion of the hidden states.

To initialize *P*_*E*_(*H*_*i*_, *H*_*j*_), we estimate it by the transition probability of initial states called from the Gaussian mixture model. For each bin *i*, we choose *H*_*i*_ to be the state *h*_*a*_ that maximizes *P* (*H*_*i*_ = *h*_*a*_|*O*_*i*_) of Gaussian mixture model. The initial transition matrix between two state *h*_*a*_ and *h*_*b*_ can be calculated as:

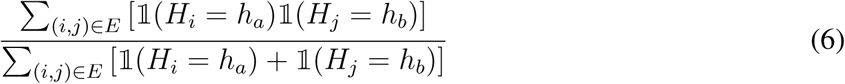

where 𝟙 is the indicator function.

We use the Expectation-Maximization (EM) algorithm to estimate the parameters in the model, including parameters to define node potential and edge potential. At iteration *t*, we assume that our estimate of model parameters from previous iteration is *θ*^*t*−1^. The goal of the EM algorithm is to maximize the expected value of the log likelihood. By using mean-fied approximation (Celeux et al., 2003; Zhang, 1992), we maximize the following *Q* function:

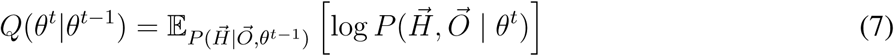

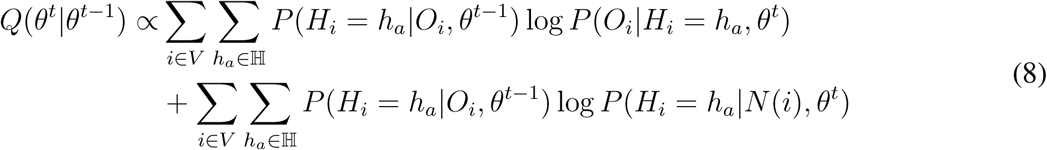

where *N* (*i*) represents the neighboring nodes of node *i*, and we can use the estimated hidden states from last iteration to approximate:

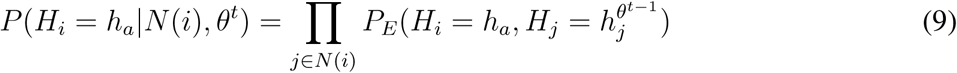

where 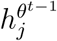 is the loopy belief propagation estimated hidden state for node *j* at iteration *t* − 1. We also have:

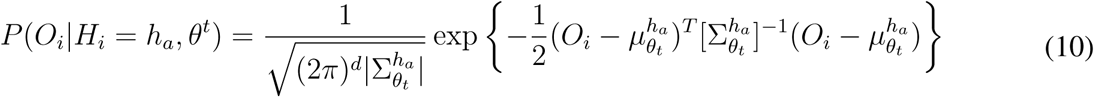

where *O*_*i*_ follows multivariate Gaussian distribution given state *H*_*i*_ = *h*_*a*_.

The first part on the right-hand side of Eqn. 8 corresponds to the node potential and the second part corresponds to the edge potential.

In the E-step, we compute the expected states of all nodes. Given the parameter estimation *θ*^*t*−1^, we calculate the posterior probability:

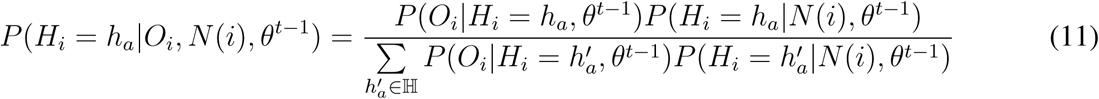

In the M-step, we use the maximum likelihood estimation (MLE) to maximize the *Q* function:

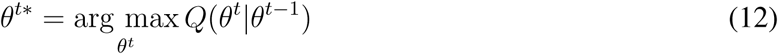

#### Loopy belief propagation for state estimation

Given the parameters and observations in the graph, the hidden state inference problem is solved by the loopy belief propagation (LBP) algorithm (Murphy et al., 1999). LBP works by passing messages among neighboring nodes in the Markov random field structure. Each node passes messages to neighboring nodes when it has received all incoming messages. The passed message from node *i* to node *j* about hidden state *H*_*j*_ = *h*_*b*_ is calculated as:

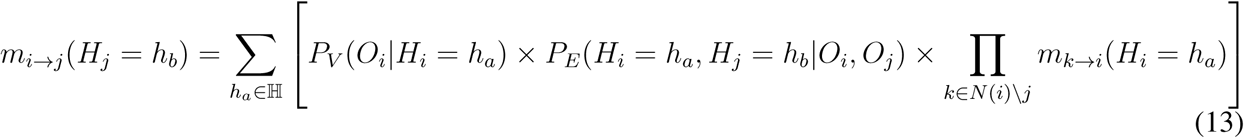

where ℍ is the set of all states, *m*_*i*→*j*_(*H*_*j*_ = *h*_*b*_) represents the message passing from node *i* to node *j* about hidden state *H*_*j*_ = *h*_*b*_. *N* (*i*) \ *j* refers to the neighbors of node *i* other than node *j*. The complete message passed between nodes should be normalized before sending. We normalize the sum of message *m*_*i*→*j*_ to be 1, i.e.,

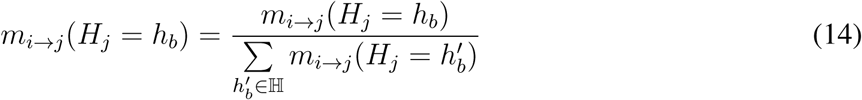

After we send messages from all nodes to their neighbors, we calculate the belief of each nodes based on the node potential and the incoming messages. The belief of node *i* with hidden state *H*_*i*_ is calculated as:

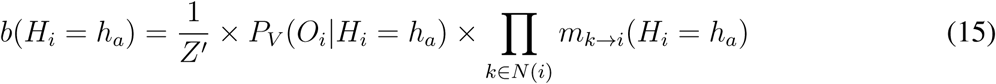

where *Z′* is the normalization constant:

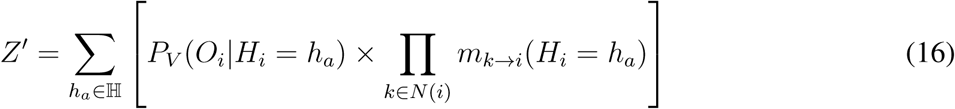

Belief is the normalized product of all incoming messages and node potentials, which approximates the marginal probability of each node. Based on belief, we can update the estimated states of each node. To do that, we simply go through all possible hidden states and choose the one with highest belief. LBP runs by iteratively passing messages among neighboring nodes and updating all messages to be sent simultaneously based on previous incoming messages. At the first iteration the initial messages are all set to 1 before they are normalized. As for the termination condition, we will stop iterations if there is no change of belief or maximum iteration number is reached. We set the maximum iteration to 500 but the method can typically terminate within 100 iterations. All computations are performed in log space to avoid numerical underflow.

#### Estimation of the number of states

To estimate the number of states, we applied the Elbow method on the input TSA-seq and DamID data based on *K*-means clustering. Specifically, we assessed the total within-cluster sum of squares as a function of the number of clusters. The total within-cluster sum of squares is calculated as:

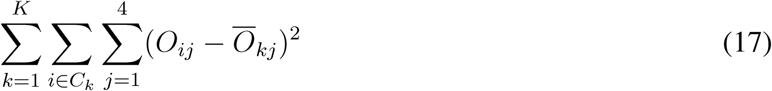

where *K* is the number of states, *C*_*k*_ is the set of cluster *k, j* refers to the 4 different input data types (TSA-seq and DamID). *Ō*_*kj*_ is the average score for data *j* in cluster *k*. We determined the appropriate cluster number *K* where additional cluster would not lead to much improvement in terms of the total within-cluster sum of squares. We found that the appropriate number of states may range from 10 to 15 (Fig. S13). However, results with different state number within the range showed minor difference. We therefore set *K*=10 in this work.

### Method for predicting DNA replication timing from SPIN states

We developed a predictive model to demonstrate that multi-fraction Repli-seq can be predicted based on SPIN states and histone marks signals. The model takes SPIN states and 11 histone modification ChIP-seq signals as input, and the 7 fraction Repli-seq score as predictive output. Here we used 25kb as window size. We then calculated the average ChIP-seq signals (*p*-value given by MACS2) with each window for 11 histone marks, H2A.Z, H3K27me3, H3K4me1, H3K4me3, H3K9ac, H3K9me3, H3K27ac, H3K36me3, H3K4me2, H3K79me2, and H3K9me1. Regions with missing values in any dataset were discarded. The discrete SPIN states were transformed into integer numbers ranging from 1 to 10, ranked by the distance from nuclear speckles (1 as Lamina state and 9 as Speckle state). The predicted Repli-seq score for each bin is a 7-dimension vector, where each dimension corresponds to a specific fraction (S1-S2, and G2).

We then utilized the random forest regressor in scikit-learn (Pedregosa et al., 2011). For the prediction model, the input contains SPIN states and histone marks averaged over 25kb bins. The performance of our model was measured by the average *R*^2^ score between real Repli-seq signal and the predicted one in all fractions. We performed a cross-validation on different chromosomes to avoid over-fitting, where we left out one chromosome as test set, and used the remaining chromosomes as training set. The process was repeated for every chromosome and then the results from each fold were averaged. To improve the predictive performance, we performed a parameter scanning for the random forest model and used the parameter set with the highest *R*^2^ score on the training set. The parameters that were tuned include the number of trees (1000), the maximum number of features in each tree (square root of the total number of features), and the maximum depth of the tree (100). The feature importance reported by the random forest regressor was used to select informative features.

We were able to achieve 0.95 *R*^2^ score on the training set and 0.923 *R*^2^ score on the testing set with consistent performance across chromosomes (Fig. S4).

## Code Availability

The source code of SPIN can be accessed at: https://github.com/ma-compbio/SPIN.

## Acknowledgement

This work was supported by the National Institutes of Health Common Fund 4D Nucleome Program grant U54DK107965 (A.S.B., B.v.S., D.M.G., and J.M.) and National Institutes of Health grant R01HG007352 (J.M.).

## Author Contributions

Conceptualization, J.M.; Methodology, Y.W. and J.M.; Software, Y.W.; Resources, L.Z., T.v.S., T.S., Y.C., D.P.H., D.M.G., B.v.S., A.S.B.; Investigation, Y.W., Y.Z., R.Z., D.M.G., B.v.S., A.S.B., and J.M.; Writing – Original Draft, Y.W., Y.Z., R.Z., and J.M.; Writing – Review & Editing, Y.W., Y.Z., R.Z., D.M.G., B.v.S., A.S.B., and J.M.; Funding Acquisition, A.S.B., B.v.S., D.M.G., and J.M.

