## Supplemental Information for "SPIN reveals genome-wide landscape of nuclear compartmentalization"

#### Supplementary Methods

##### Multi-fraction Repli-seq data generation

Exponentially growing K562 cells (ATCC CCL-243) cultured in IMDM-10% FBS were pulse-labeled with 200 $\mu$M BrdU for 60 minutes, then fixed in 70% ethanol and stained with propidium iodide as previously described for sorting by DNA content (Marchal et al., 2018). FACS sorting windows were defined as previously published (Hansen et al., 2010). Briefly, the area between G1 and G2 peaks were divided into 6 windows evenly (S1-S6). Another window (G1) was added immediately on the left of S1 to collect non-replicating cells. At least 40,000 cells were collected from each fraction, then using 40,000 cells each, sequencing libraries were made. From G1 fraction, the total genome library was made as the control. From S1-G2 fraction, libraries that represent nascent DNA were made by performing anti-BrdU immunoprecipitation as described (Marchal et al., 2018). These libraries were sequenced on HiSeq 2500 with 50SE chemistry aiming to obtain 20-40 million reads each.

##### Simulation data generation

To evaluate the performance of SPIN, we generated simulation datasets and directly compared the results with baseline methods such as $K$-means and Gaussian HMM. For simulation, we first generated a graph on chromosome 1 at 25kb resolution where edges in the graph represent the significant Hi-C interactions. We kept the ratio between the local interactions (<1Mb) and long-range interactions (>1Mb) as 3:1, similar to the ratio derived from the real data. Given the graph structure, we used Gibbs sampling to generate a series of simulated states and the corresponding TSA-seq and DamID signals. First, we initialized the graph with random state labels. Then the Gibbs sampling sampled on each node in the graph, one at a time. Gibbs sampling can take advantage of the conditional independence properties of the Markov random field. Thus, when sampling the state $H_i$ of node $i$, we only need to consider its direct neighboring nodes $N(i)$. The state $H_i$ was sampled from distribution: $P(H_i|\{H_j|_{j\in N(i)}\})$. We iteratively updated node state labels until convergence (no changes in state labels). Given the simulated state, TSA-seq and DamID signals on each node were sampled using the distribution learned from the real data and then smoothed. We applied SPIN to estimate the hidden states based on TSA-seq, DamID, and the graph structure as input. As a comparison, we also estimated the states using $K$-means and Gaussian HMM, which only used TSA-seq and DamID signals as input. We evaluated the performance of different method. In addition, we calculated the rand index to evaluate the model performance. Fig. S14 shows an example of the simulated data.

##### Considering structural variations in K562

Since K562 is a cancer cell line, we also took large structural variants (SVs) into account. For example, if genomic region $s$ and $t$ are adjacent in cancer genome but not in the normal genome, we would add an edge $(s, t)$ in the HMRF graph structure. We applied our recently developed SV caller Weaver (Li et al., 2016) to generate SVs in K562 based on the whole-genome sequencing (WGS) data from the CCLE project (Barretina et al., 2012). Weaver identified 121 large-scale (allele-specific) SVs in K562 (Table S2). 64% of Weaver identified SVs were also reported in Dixon et al. (2018). We then integrated these SVs into our HMRF graph construction by adding/deleting edges (see below). Overall, 46 edges due to SVs were added and 29 edges were removed. First, we assigned each breakpoint of SVs to the closest boundary of non-overlapping 25kb genomic bins. We modified the original graph with SV information using the following steps:

1. We call two breakpoints involved in a SV $X$ and $Y$, and the genomic bins upstream/downstream of breakpoint $X$ are referred to as $X_L$ and $X_R$, respectively. Similarly, the genomic bins upstream/downstream of breakpoint $Y$ are referred to as $Y_L$ and $Y_R$.
2. We count the copy number of each bin on two alleles separately (as part of the output from Weaver). We denote the two alleles as A and B, respectively. For example, the copy number of bin $X_L$ on allele A is notated as $C(X_L^{(A)})$ and the copy number on allele B is $C(X_L^{(B)})$.
3. Deletion of edges: for breakpoint $X$, we check the connectivity of the chromosome on each allele based on copy number.

$$Con(X^{(A)}) = \begin{cases} 0, & \text{if } C(X_L^{(A)})) = 0 \\ 0, & \text{if } C(X_R^{(A)})) = 0 \\ 1, & \text{otherwise} \end{cases} \tag{18}$$

   where $Con(X^{(A)})$ stands for the connectivity of allele $A$ at breakpoint $X$. $Con(X^{(A)}) = 0$ indicates that all copies of allele A are completely disconnected at breakpoint $X$. $Con(X^{(A)}) = 1$ indicates that at lease one copy of the chromosome is connected at breakpoint $X$. The genome is disconnected at breakpoint $X$ when both alleles are broken. In other words, we will delete edge $(X_L, X_R)$ in the original graph when $Con(X^{(A)}) = 0$ and $Con(X^{(B)}) = 0$. Otherwise, we will keep the original edge because at least one copy of the chromosome is connected near breakpoint $X$. We do the same for breakpoint $Y$.
4. Adding edges. For breakpoints $X$ and $Y$, we calculate the difference in copy numbers across break points. For example, the difference of copy number for breakpoint $X$ is defined as:

$$D(X) = C(X_L^{(A)}) - C(X_R^{(A)}) + C(X_L^{(B)}) - C(X_R^{(B)}) \tag{19}$$

   If $D(X) = 0$, we do not expect SVs at breakpoint $X$. Otherwise, we add edges to the graph to reflect the SV. Typically we only observe change of copy number on one of the allele. The orientation of edges to add can be determined by $D(X)$ and $D(Y)$.

#### Collecting results from other chromatin state annotation methods

We compared the SPIN states with other chromatin state annotations, including Hi-C subcompartments, chromatin state annotations from ChromHMM (Ernst and Kellis, 2012) as well as Segway (Hoffman et al., 2012a) and Segway-GBR (Libbrecht et al., 2015).

For Hi-C subcompartments, we used the SNIPER Hi-C subcompartments in K562 (Xiong and Ma, 2019), although the trend with SPIN state comparison is largely consistent with the original Hi-C subcompartment definitions from GM12878 (Rao et al., 2014). We focused on the five primary Hi-C subcompartments, i.e., A1, A2, B1, B2 and B3.

To compare SPIN states with Hi-C subcompartments, we calculated the genome-wide coverage fold change of each SPIN states over each Hi-C subcompartment. For each Hi-C subcompartment, we calculated the total length of regions covered by different SPIN states, and normalized it by the expected length, which was calculated by the genome-wide coverage of SPIN states (as shown in Fig. 1c). We then defined the ratio between the observed length and the expected length as the fold change score of the SPIN states. If one SPIN state is uniformly distributed along the genome, its fold change score for each Hi-C subcompartments should be 1. Higher fold change score indicates certain SPIN states are more likely to be found in a given Hi-C subcompartment.

We used the SNIPER Hi-C subcompartment calls in all 9 human cell lines: K562, GM12878, HAP1, HeLa, HMEC, HSPC, HUVEC, IMR90, and TCell from Xiong and Ma (2019) to estimate the constitutive level of subcompartment annotations. We calculated entropy as a metric for the level of conservation of Hi-C subcompartments across different cell types. Within each 25kb genomic bin, we have the Hi-C subcompartment calling in 9 cell lines and, for each subcompartment $i$, we use $l_i$ to denote the total genomic length across 9 cell lines. The entropy was thus calculated as:

$$E = -\sum_{i \in \{A1, A2, B1, B2, B3\}} \left(\frac{l_i}{L}\right) \log\left(\frac{l_i}{L}\right) \tag{20}$$

where $L$ is the sum of genomic length for all subcompartments in that genomic bin across cell types.

We downloaded ChromHMM and Segway chromatin state annotations in K562 from the ENCODE Project. In K562, there are 25 ChromHMM states and 25 Segway states identified from histone modifications, transcription factor binding, and open chromatin data. We also downloaded the combined chromatin state definition in K562, which is a consensus merged from the results of ChromHMM and Segway (Hoffman et al., 2012b). There are seven states in the combined segmentation. We calculated the total length of the overlapping regions between ChromHMM/Segway/Combined and the SPIN states, and then normalized it by the expected length. The expected length was calculated by the genome-wide coverage of the SPIN states. We then calculated the fold enrichment of chromatin states from ChromHMM, Segway, and Combined, respectively, over each SPIN state. For Segway-GBR chromatin domain annotations, we used the original Segway-GBR annotations in Libbrecht et al. (2015) for GM12878 and IMR90 cell lines. For fair method comparison with Segway-GBR, instead of using TSA-seq and DamID as input, we ran SPIN using the same input with Segway-GBR in IMR90 and GM12878 cell lines. The input data include 11 histone modification (H2A.Z, H3K27ac, H3K27me3, H3K36me3, H3K4me1, H3K4me2, H3K4me3, H3K79me2, H3K9ac, H3K9me3, H4K20me1), and DNase-seq. The histone modification and DNase-seq signals were averaged on 25kb bins and reduced dimensions using PCA. The first 5 principal components were used as input for SPIN. For Hi-C, we used H1 and IMR90 Hi-C data from Dixon et al. (2012) (same as what Libbrecht et al. (2015) used). We downloaded 6-fraction Repli-seq in IMR90 and GM12878 from the ENCODE project to evaluate the performance of these two different methods.

#### Methods for sequence feature analysis in SPIN states

To identify sequence features associated with SPIN states, we focused on quantifying of *k*-mers, transposable elements (TEs), and TF motifs on SPIN states. For *k*-mers, we counted 6-mer occurrences on 25kb genomic windows using Jellyfish (Marçais and Kingsford, 2011). Then we used t-SNE (Maaten and Hinton, 2008) to visualize all possible 6-mers, where each dot represents a 25kb genomic bin with its SPIN states shown in colors (Fig. S11).

For transposable elements (TEs), we downloaded the annotations of TE families from the UCSC Genome Browser (Hinrichs et al., 2006). The 16 TE families that we used are srpRNA, scRNA, snRNA, Alu, MIR, L1, L2, CR1, ERVL, ERV1, Gypsy, ERVK, LTR, DNA, hAT, and PiggyBac. We further grouped the 16 transposable elements families into 5 classes based on their sequence properties (RNA,

SINE, LINE, LTR, and DNA). We counted the density of each TE family on different SPIN states and compared to the genome-wide average.

For known TF motifs, we used the 730 human TF motifs from CIS-BP (Weirauch et al., 2014) and scanned on the open chromatin regions of the whole genome for matches of each motif. The open chromatin regions were defined from DNase-seq peaks in K562 from the ENCODE project (accession number: ENCSR921NMD). Peak calling of DNase-seq was done by the ENCODE standard pipeline. We used FIMO with $p$-value cutoff $10^{-5}$ for motif matching. The known TF motifs were further grouped into 12 classes based on the annotation from CIS-BP.

To search for motifs *de novo* on different SPIN states, we used MEME (Bailey et al., 2009). We used MEME (Ma et al., 2014) for motif discovery and motif enrichment analysis. On each SPIN state, we extracted randomly sampled DNA sequences on that SPIN state as the primary sequence set and randomly sampled DNA sequences of the whole genome as the control sequence set. We set the length of motifs to be between 6 and 50 nt. We then used FIMO to find the distribution of discovered motifs. We also used Tomtom to match identified motifs to known TF motifs. The known motifs used here were the human TF motifs from HOCOMOCO v11.

### Supplementary Results

#### Comparison with other genome segmentation methods

We compared SPIN to Segway (Hoffman et al., 2012a) and ChromHMM (Ernst and Kellis, 2012) results in K562 based on histone modification data (Fig. S15). We also included combined annotations of Segway and ChromHMM from ENCODE. The fold enrichment of each state is calculated. Overall, we found similar results by comparing to Segway and ChromHMM (Fig. S15). Most enhancer (Enh-), promoter (Prom), and gene related states (Gen, Tss) have stronger presence in the Speckle and Interior_Act states. Quiescent state (Quies) has more involvement in the Lamina state and the repressed polyComb state (Repr) is enriched in the Interior_Repr2 state. For combined annotation of Segway and ChromHMM, we found that TSS, T (transcribed), E (Enhancer), WE (weak enhancer), and PF (promoter flanking) all have strong presence in the Speckle and Interior_Act states. R (Repressed) state is enriched in the Lamina Interior_Repr and Near_Lm states. These results are consistent with our observations that spatial localization is highly correlated with chromatin states and transcription. Most gene rich and active chromatin regions are in the Speckle and Interior_Act states. Polycomb/facultative repressed regions are in the Interior_Repr state. Heterochromatin and low transcription activity regions are in the Near_Lm and Lamina states.

We also compared SPIN with Segway-GBR (Libbrecht et al., 2015), which uses graph-based regularization (GBR) to integrate pairwise chromatin conformation data for genome segmentation (Fig. S16). In order to compare the two methods, we used the same input data in IMR90 and GM12878 from Libbrecht et al. (2015) to run SPIN and set the number of states to be 5. We used 6-fraction Repli-seq from ENCODE as a way to evaluate the segmentation performance. We expect that a better segmentation will better stratify Repli-seq data. We found that the states produced by SPIN have clearer Repli-seq stratification as compared to Segway-GBR. SPIN results can distinguish G1 from S1 and G2 from S4 fraction. In contrast, the Repli-seq patterns are similar between SPC and BRD, QUI and CON in the Segway-GBR results.

#### Simulation study to evaluate the performance of SPIN

To assess the advantage of using SPIN to infer compartmentalization patterns, we compared the results from SPIN to the baseline methods $K$-means and Gaussian HMM on the simulated data. We simulated TSA-Seq SON, TSA-Seq LaminB, DamID LaminB, and DamID Nucleolus scores with 25kb window size on chromosome 1 (Supplementary Methods, Fig. S14). In order to generate TSA-Seq and DamID scores similar to the real data, we used 5 major SPIN states call in K562 (Speckle, Interior_Act, Interior_Repr, Near_Lm, and Lamina) as ground truth and used multivariate Gaussian distribution of each state to generate new TSA-Seq/DamID scores. Then the TSA-Seq/DamID is smoothed with using Hanning window of length 21, and used as input for SPIN, $K$-means and Gaussian HMM. For fair comparison, we set the number of states to be 5 for $K$-means, Gaussian HMM, and SPIN. We evaluated the performance comparing prediction accuracy of each state (Table S4). We found that SPIN outperforms both $K$-means and Gaussian HMM on all states. SPIN states have high prediction accuracy on the Speckle state (96%), Interior_Act (91%), and Lamina states (86%), with notable improvement as compared to Gaussian HMM. For Near_Lm and Interior_Repr, SPIN still perform much better than the other two methods. The simulation results suggest that SPIN can identify compartmentalization patterns much more accurately than baseline $K$-means and Gaussian HMM.

#### TAD-TAD interactions in SPIN states

Recently, Paulsen et al. (2019) showed that long-range TAD-TAD interactions form constitutive TAD cliques and the formation of TAD cliques stabilizes heterochromatin at the nuclear lamina. However, it is unclear how other large TAD cliques are spatially localized and contribute to genome compartmentalization. Here we analyzed TAD-TAD interactions in K562 and compared them to the SPIN states (Fig. S6). We first used the DI method to call TADs, and we calculated the median of O/E Hi-C interaction between each pair of TADs. For pairs with TADs with median O/E >2, we found that 55.6% are between the same SPIN states, and 31.2% are between the similar SPIN states (e.g., between Interior_Act states). The Lamina state has the most self TAD-TAD interaction with over 60% of its interactions are with another TAD in the Lamina state. This is consistent with observations in Paulsen et al. (2019). Speckle and Interior_Act3 TADs also tend to self-interact (over 40%). If we separate the TADs into two groups based on their SPIN states, interior (Speckle, Interior_Act, Interior_Repr) and periphery (Near_Lm, Lamina_Like, Lamina), we found that there is only <1% interactions are between interior and periphery TADs. Our results support the hypothesis that TADs form spatially separated cliques. We found that TAD cliques at the Lamina states are the strongest and there is little TAD-TAD interactions between a pair of TADs that are localized separately at the interior and periphery.

#### SPIN states harbor distinct preferences in DNA binding proteins

We explored the DNA binding proteins in different SPIN states based on 381 ChIP-seq datasets in K562 from the ENCODE project. We utilized the conservative peaks provided by ENCODE if available and called peaks using MACS2 (Zhang et al., 2008) if the peaks were not available. For ChIP-seq dataset with replicates, we kept peak regions shared in both replicates. We calculated the ratio of observed and expected length of ChIP-seq peaks in different SPIN states (i.e., the fold enrichment of ChIP-seq peaks in each SPIN state). To find proteins with similar patterns of occupancy in different SPIN states, we performed hierarchical clustering of fold enrichment score for peaks that led to three clusters (Clusters 1-3) (Fig. S8a). We found that the majority of DNA binding proteins (314; >80%) preferentially bind to the Speckle and Interior_Act states (Cluster 2). Examples include CTCF, SMC3, SP1, and STAT1. 22 of them (including BRCA1, SRSF9, RBM17, and NONO) have ChIP-seq peaks enriched in the Lamina and Near_Lm2 states (Cluster 1). There are also 46 DNA binding proteins (including AGO1, U2AF1, KLF16, and FOXA1) that show preference in binding to Interior_Act2 and Interior_Act3 states (Cluster 3). This clustering analysis demonstrates that even though most of the proteins preferentially bind to transcriptionally active SPIN states, there are many that bind to more target genes in the transcriptionally less active SPIN states (e.g., Interior_Act2 and 3). We also showed that the majority of RBPs have binding sites in the Speckle state (Fig. S17).

We then asked whether the target genes of the DNA binding proteins in different spatial clusters based on the SPIN states exhibit functional enrichment. For each clusters, we merged all ChIP-seq peaks called by MACS2, and used the merged list of peaks as input for GREAT (McLean et al., 2010). As shown in Fig. S8a and Table S5, the most enriched GO terms in different clusters vary, suggesting different roles of these DNA binding proteins and their target genes in the entire transcriptional regulatory network.

#### SPIN facilitates the identification of sequence features for nuclear compartmentalization

We asked what sequence features are enriched in different SPIN states. We analyzed the distribution all known human TF motifs (and then grouped by families) on different SPIN states using CIS-BP motifs (Weirauch et al., 2014). We found that some motif families (POU, Forkhead, Hemedomain, SOX, and Rel) are highly enriched in the Lamina and Near_lm2 states ($p$-value $<$2.2E-16) (Fig. S8b). Motif families such as ETS, bHLH, Nuclear receptor, and C2H2 have more presence in the Speckle state ($p$-value $<$2.2E-16). These enriched TF motifs suggest that the sequence motifs and also the corresponding TF families may also play roles in cooperatively modulating global spatial localization.

We next sought to identify TF binding motifs *de novo* in different SPIN states. Here we used DNA sequences of each SPIN state as input for motif discovery using MEME (Bailey et al., 2009) (Fig. S9). We identified 15 motifs enriched in different SPIN states (Fig. S9), and matched them to known human motifs (Fig. S10). Specifically, for the Speckle state, we identified three motifs matched to the known motifs of MAZ, ZN467, VEZF1, WT1, SP1, SP2 and KLF5 with $p$-value $<$10E-5 (Fig. S9). These motifs belong to the C2H2 Zinc Finger families that are enriched in the Speckle state (Fig. S8b) and have been shown to play roles in transcriptional activation (Chen et al., 2004)..

Earlier studies have shown that transposable elements (TEs) correlate with nuclear localization (Guelen et al., 2008; Rhind and Gilbert, 2013; Yang et al., 2018). We assessed the association of TE families in each SPIN state based on repeat families with over 1000 occurrence on the whole genome that were subsequently grouped into five groups (see Supplementary Methods). For each TE family, we calculated the fold enrichment of repeat counts on each SPIN state compared to expected (Fig. S8c). We found that different TE families have different preferred spatial localization. All three small RNAs (srpRNA, scRNA, snRNA) are highly enriched in the Speckle and Interior_Act states (fold enrichment $>$1.5, $p$-value$<$2.2E-16), and depleted in other states. For the SINE family, Alu is mostly abundant in the Speckle state (fold enrichment 2.6) followed by the Interior_Act states (fold enrichment $>$1.5). There is also enrichment of L1 elements in the Lamina and Interior_Repr2 states ($p$-value=3.3E-6), consistent with previous report that L1 elements have higher density on LADs (Guelen et al., 2008). In addition, ERVK has significant abundance in the Lamina_Like and Interior_Repr1 states ($p$-value$<$2.2E-16). Taken together, these results further demonstrate the difference in sequence features associated with different SPIN states, suggesting their potential roles in modulating nuclear compartmentalization globally.

##### Analysis of three consecutive SPIN states that reflect trajectory

We sought to ask what type of chromatin trajectories that the SPIN states can reveal. We used three consecutive SPIN states to describe the spatial arrangement of chromatin trajectory in the nucleus (Fig. S12). For example, a three consecutive Lamina–Near_Lm–Speckle region suggests a trajectory of a chromatin fiber at nuclear lamina on one end and at nuclear speckle on the other end. For each genomic region of a SPIN state, we counted its adjacent SPIN states and merge symmetrical three consecutive patterns. We ignored all genomic fragments with size smaller than 100kb. We found that all Speckle states are flanked by Interior_Act1. Similarly, almost all Lamina states are flanked by Near_Lm2. We found that for genomic regions with the same SPIN state but different three connective patterns, the histone modification and Repli-seq could be different. For example, for Interior_Act states, the histone modification and Repli-Seq signal can be different depending on whether adjacent states include the Interior_Repr or Near_Lm states. The average signal of active histone marks is 1.5 times higher if Interior_Act is not adjacent to the Interior_Repr states. These results strongly suggest that the trajectories of chromatin (or the context of SPIN state) have important implications on the functional roles of the chromatin.

#### Supplementary Figures and Tables

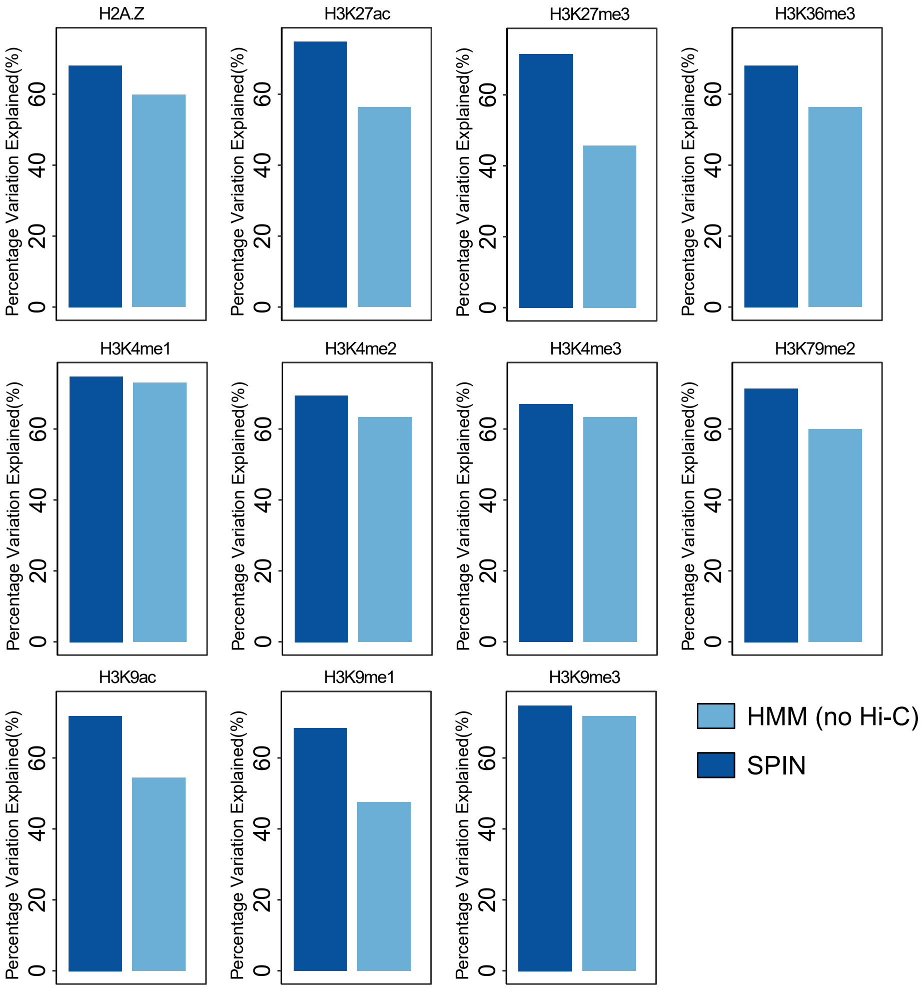


**Figure S1:** Percentage of variance for histone modification explained by the SPIN states as compared to the baseline HMM model. The HMM uses the same input (TSA-seq and DamID data) with the SPIN model except for Hi-C, and outputs the same number of states.

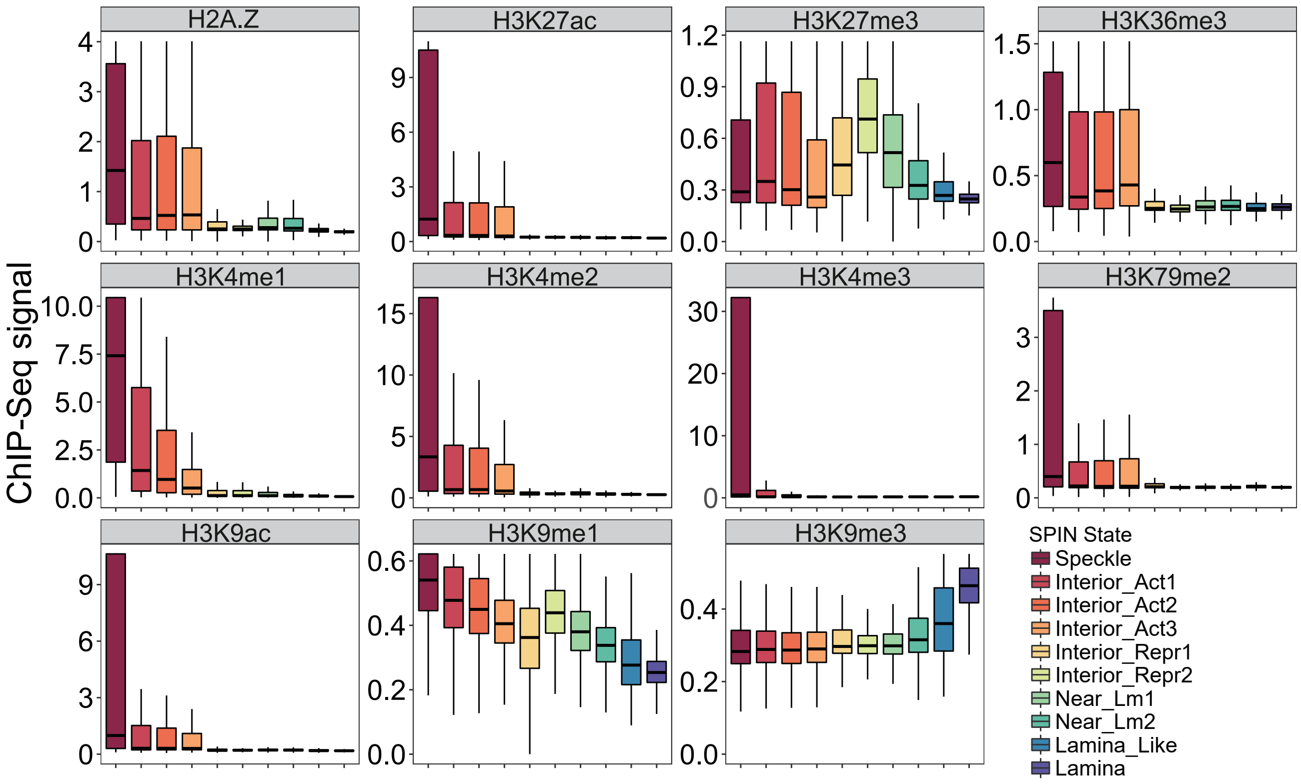


**Figure S2:** Distribution of histone modification signals on different SPIN states. 11 histone modification ChIP-seq data in K562 are shown: H2A.Z, H3K4me1, H3K9me1, H3K9me3, H3K27me3, H3K9ac, H3K36me3, H3K4me2, H3K27ac, and H3K79me2. The X-axis shows the histone modification ChIP-seq signal $p$-value calculated by MACS2.

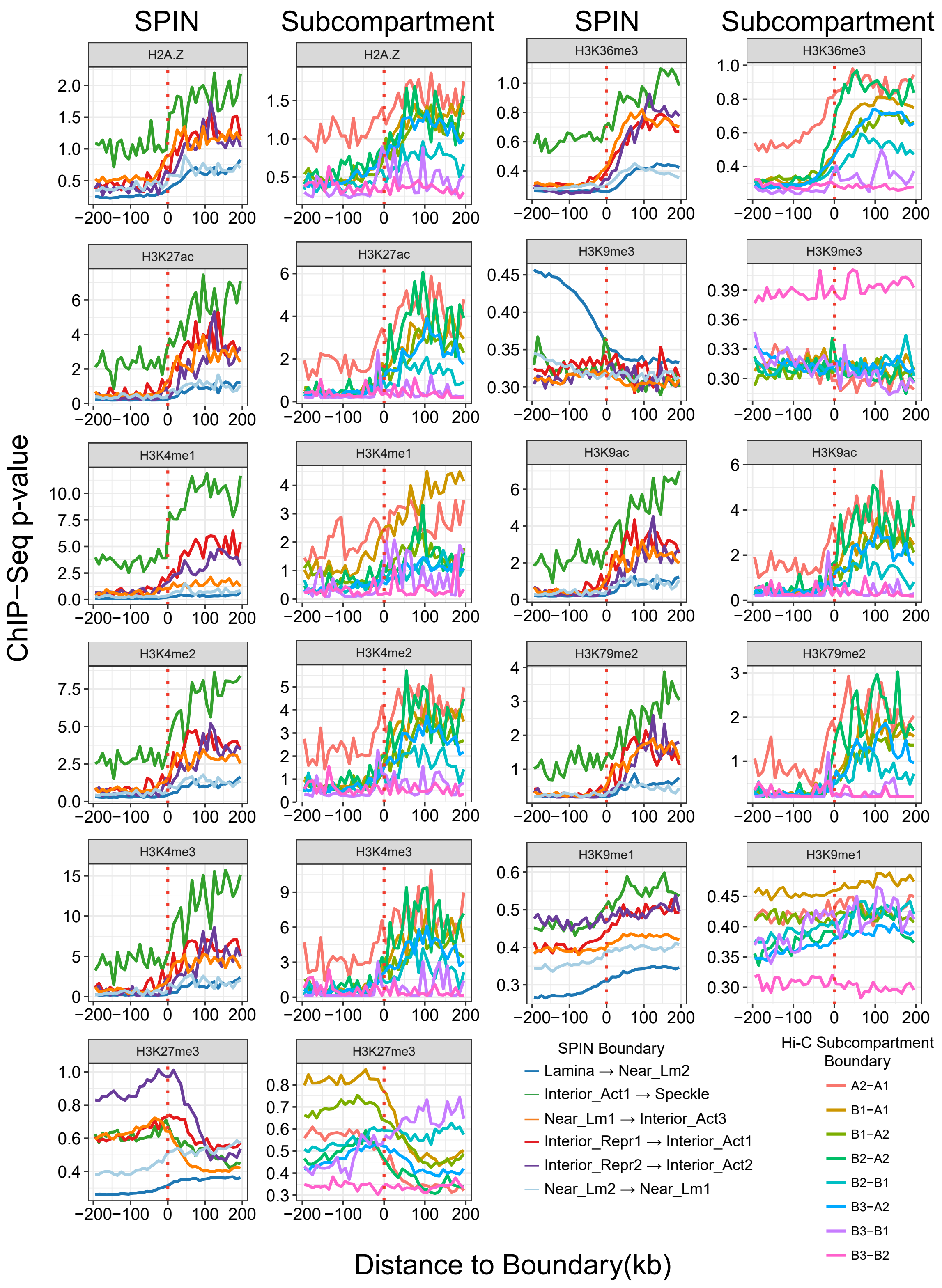


**Figure S3:** Histone modification signals at the SPIN states boundaries compared with the Hi-C subcompartment boundaries in K562 (shown side by side). Histone marks here include: H2A.Z, H3K27ac, H3K27me3, H3K36me3, H3K4me1, H3K4me2, H3K4me3, H3K79me2, H3K9ac, H3K9me1, and H3K9me3. Hi-C subcompartment annotations in K562 are from SNIPER (Xiong and Ma, 2019). Histone mark signals at most common SPIN and Hi-C subcompartment boundaries are shown. Y-axis shows ChIP-seq $p$-values for different histone marks. X-axis shows distance to state boundaries.

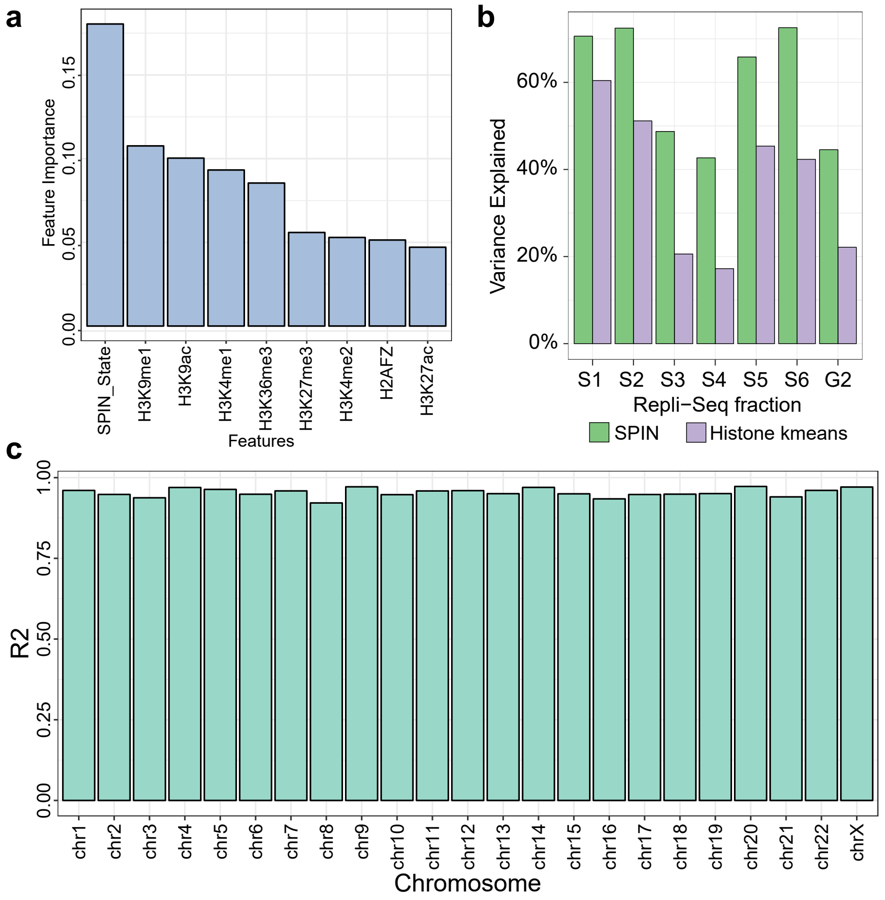


**Figure S4:** Evaluation of the model to predict Repli-seq from the SPIN states. **a.** Feature importance of the random forest model to predict Repli-seq signals. **b.** Variance of Repli-seq data explained by the SPIN states and histone marks. Histone marks are categorized using K-means clustering (K=10). **c.** Cross-validation result of predicting Repli-seq. X-axis shows which chromosome is used as test set and the remaining chromosomes are used as training set. $R^2$ score of prediction on the test set is shown in Y-axis.

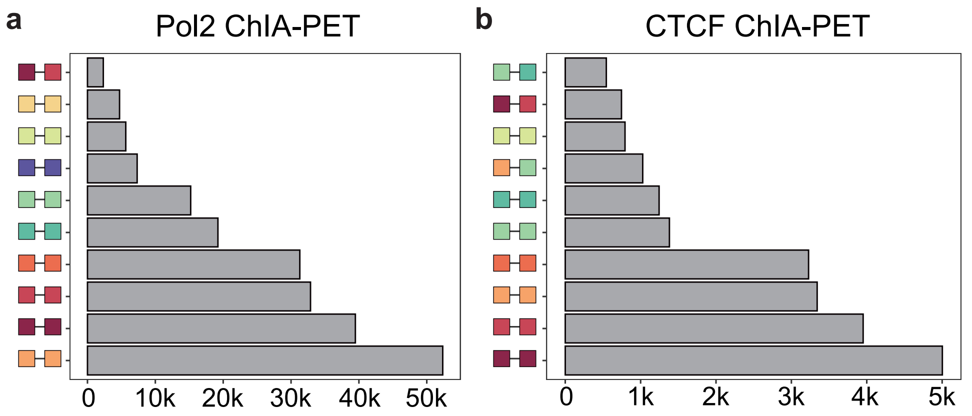


**Figure S5:** SPIN states of ChIA-PET significant interaction pairs. Only ChIA-PET pairs with distance >25kb are considered. X-axis shows the number of ChIA-PET interactions between 2 specific SPIN states. **a.** Pol2-mediated ChIA-PET loops in K562. **b.** CTCF-mediated ChIA-PET loops in K562.

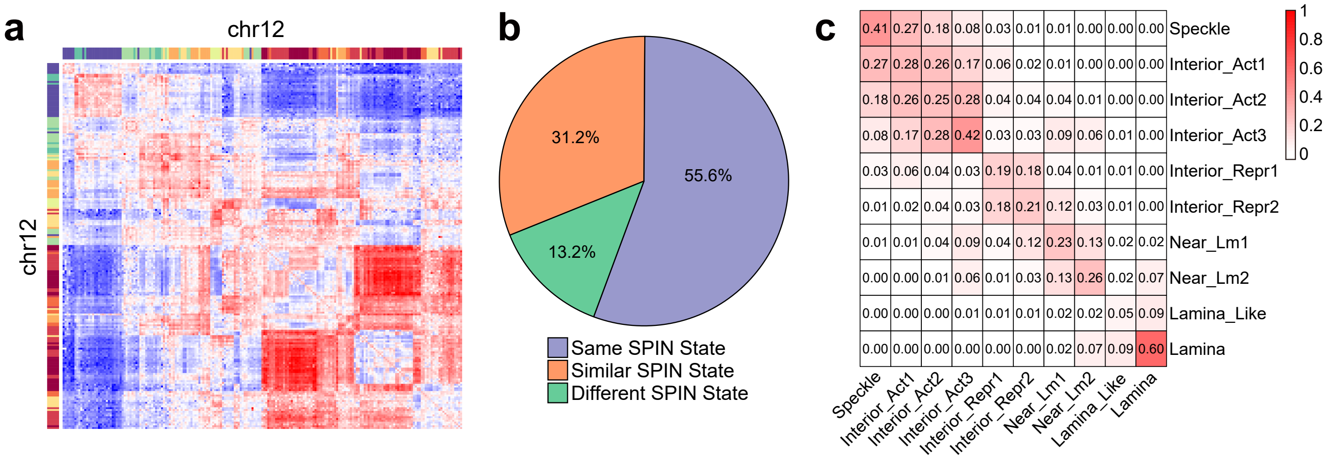


**Figure S6:** TAD-TAD interactions among SPIN states. We calculated the median O/E interactions between TADs in K562 (called from the directionality index (DI) method). **a.** Hi-C O/E matrix of TAD-TAD interactions on chr12. **b.** Pie chart showing the percentages of SPIN states among TAD-TAD interactions. **c.** Confusion matrix of TAD-TAD interactions between SPIN states.

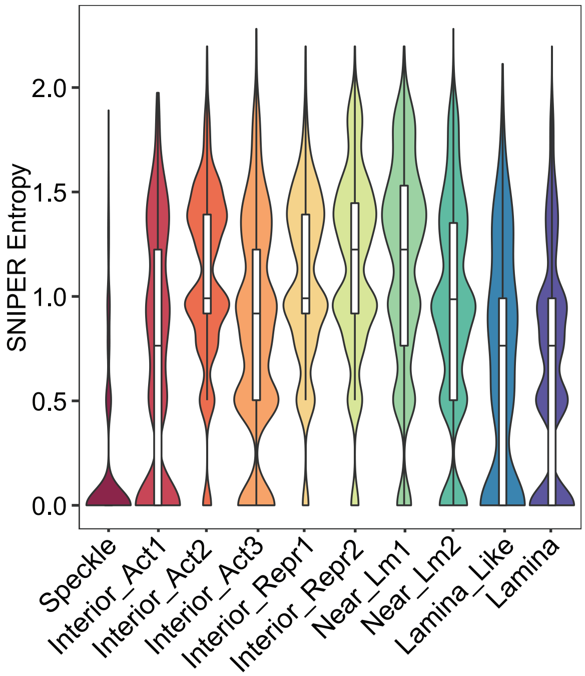


**Figure S7:** Violin plot of the entropy for SNIPER Hi-C subcompartments across cell types for different SPIN states (labeled by color). Besides violin plot, the corresponding boxplots are also shown. The entropy is calculated based on SNIPER subcompartment in 9 human cell lines (see Methods).

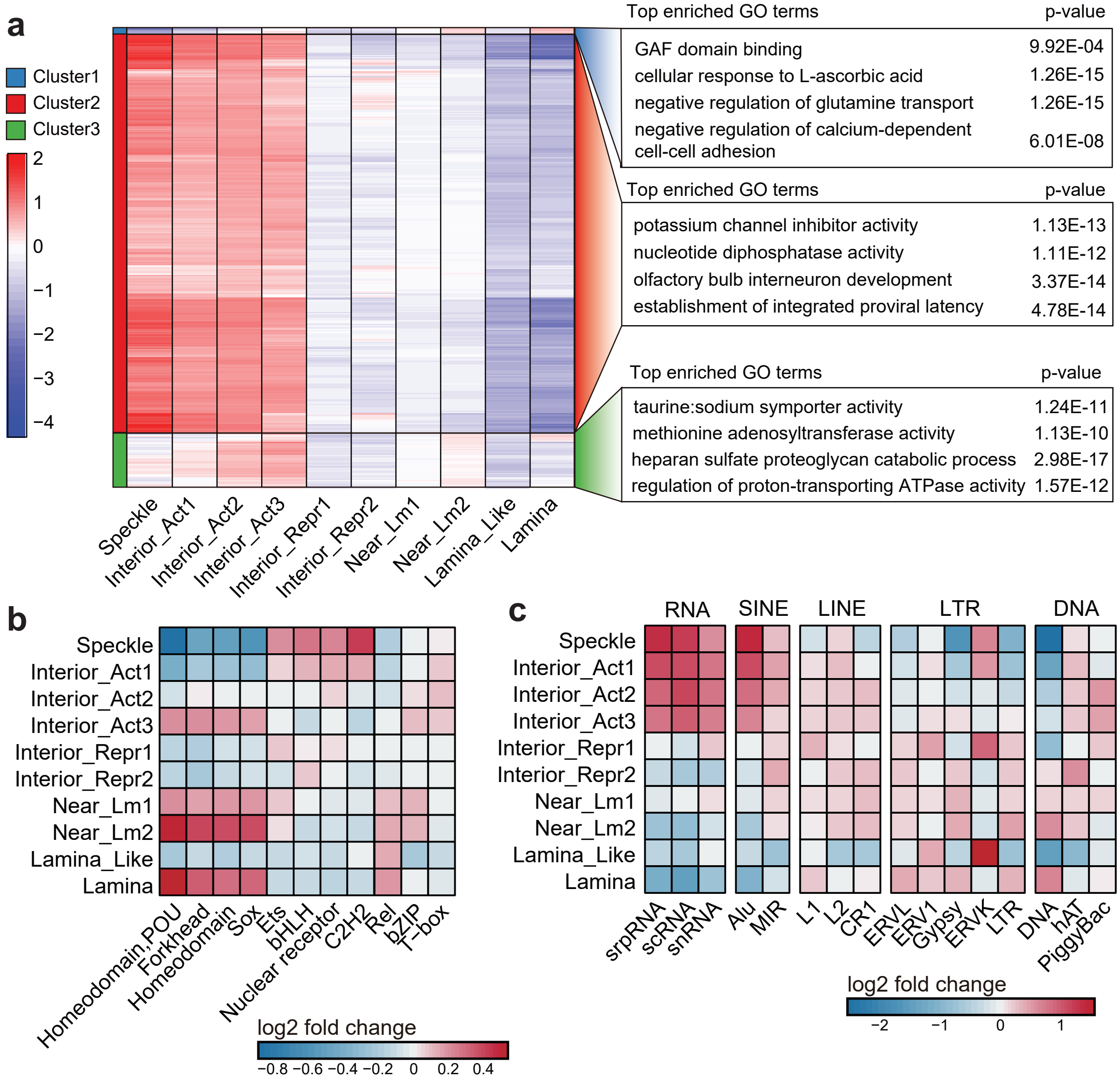


**Figure S8:** SPIN states stratify transcriptional regulation and reveal potential sequence features modulating compartmentalization. **a.** Left panel: Heatmap showing ChIP-seq peak enrichment in different SPIN states. 381 ChIP-Seq data sets are used and grouped into 3 clusters. Right panel: GO term enrichment of target genes in each cluster. **b.** Heatmap showing different transcription factor motif families enriched in each SPIN state. Motif families annotations are collected from CIS-BP. log2 fold enrichment of each motif family on SPIN states is shown. **c.** Heatmap showing different repeat families enriched in each SPIN state. Repeat families annotations are collected from the UCSC Genome Browser. log2 fold enrichment of each repeat family on SPIN states is shown.

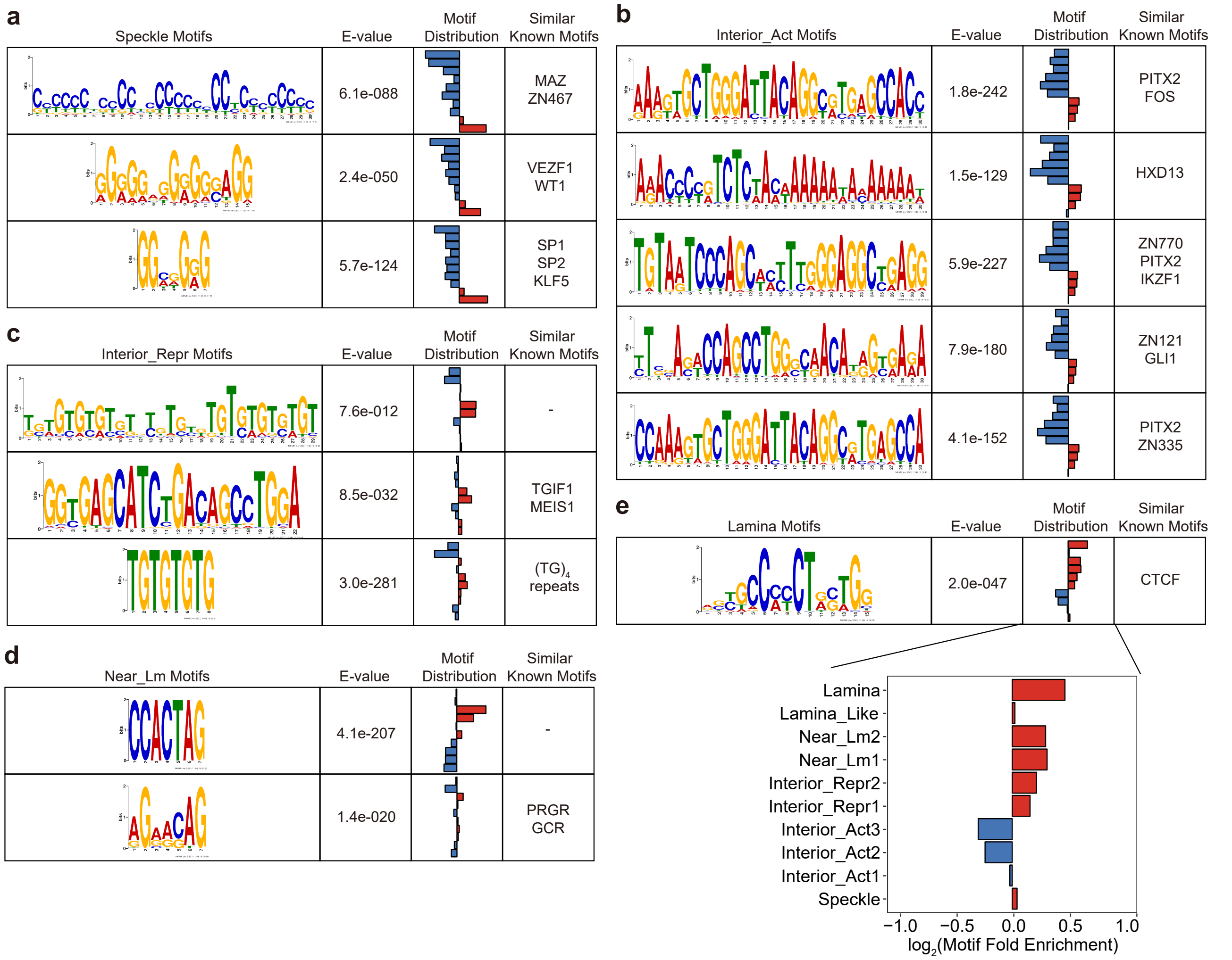


**Figure S9:** *De novo* motif discovery on different SPIN states: **a.** Speckle; **b.** Interior_Act; **c.** Interior_Repr; **d.** Near_Lm; and **e.** Lamina. We use MEME to perform motif discovery on each SPIN state. For each motif, the fold enrichment on each SPIN state is shown along with E-values. The *de novo* discovered motifs are matched to known motifs using TomTom.

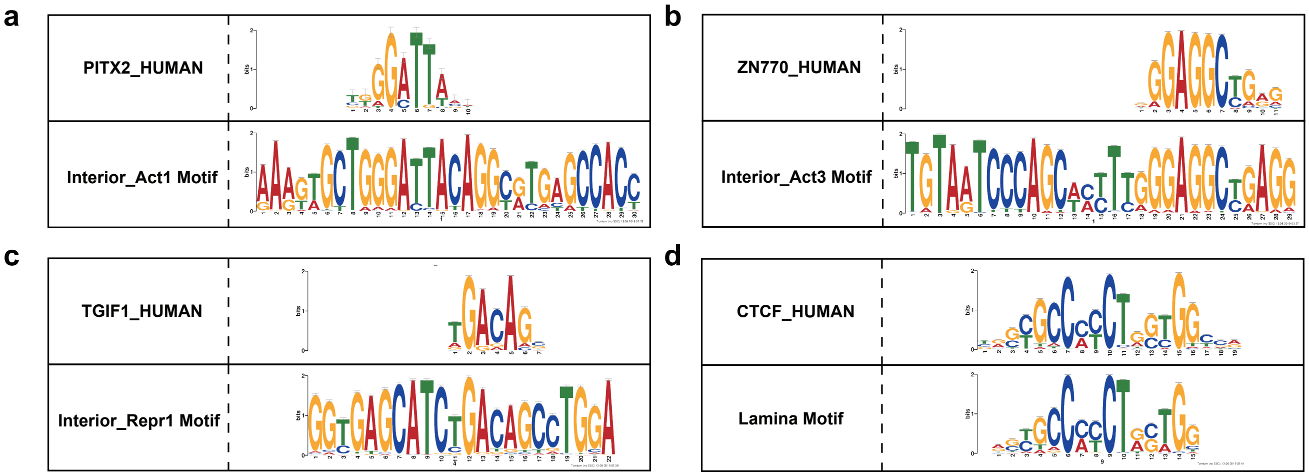


**Figure S10:** Match *de novo* discovered motifs to known motifs. Motifs discovered are compared to known motifs from JASPAR by using Tomtom from the MEME Suite. **a.** Motif enriched in the Interior_Act1 state compared with the PITX2 motif. **b.** Motif enriched in the Interior_Act3 state compared with the ZN770 motif. **c.** Motif enriched in the Interior_Repr1 state compared with the TGIF1 motif. **d.** Motif enriched in the Lamina state compared with the CTCF motif.

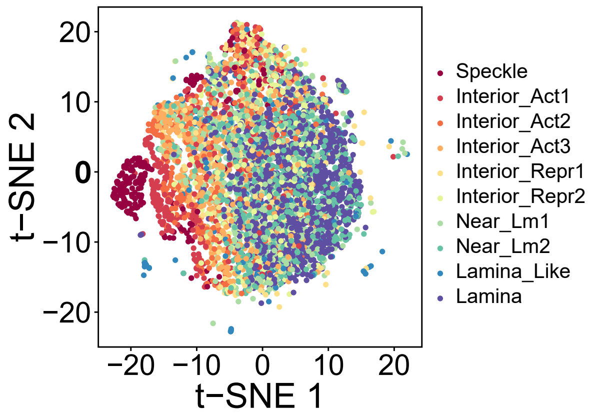


**Figure S11:** Visualization of 6-mer occurrences in sequences within different SPIN states. The 6-mer occurrences are normalized by average GC content in each SPIN state. Here we project the normalized 6-mer occurrences onto 2D plane using t-Distributed Stochastic Neighbor Embedding (t-SNE). Each dot is a 25kb bin and the color represents different SPIN states.

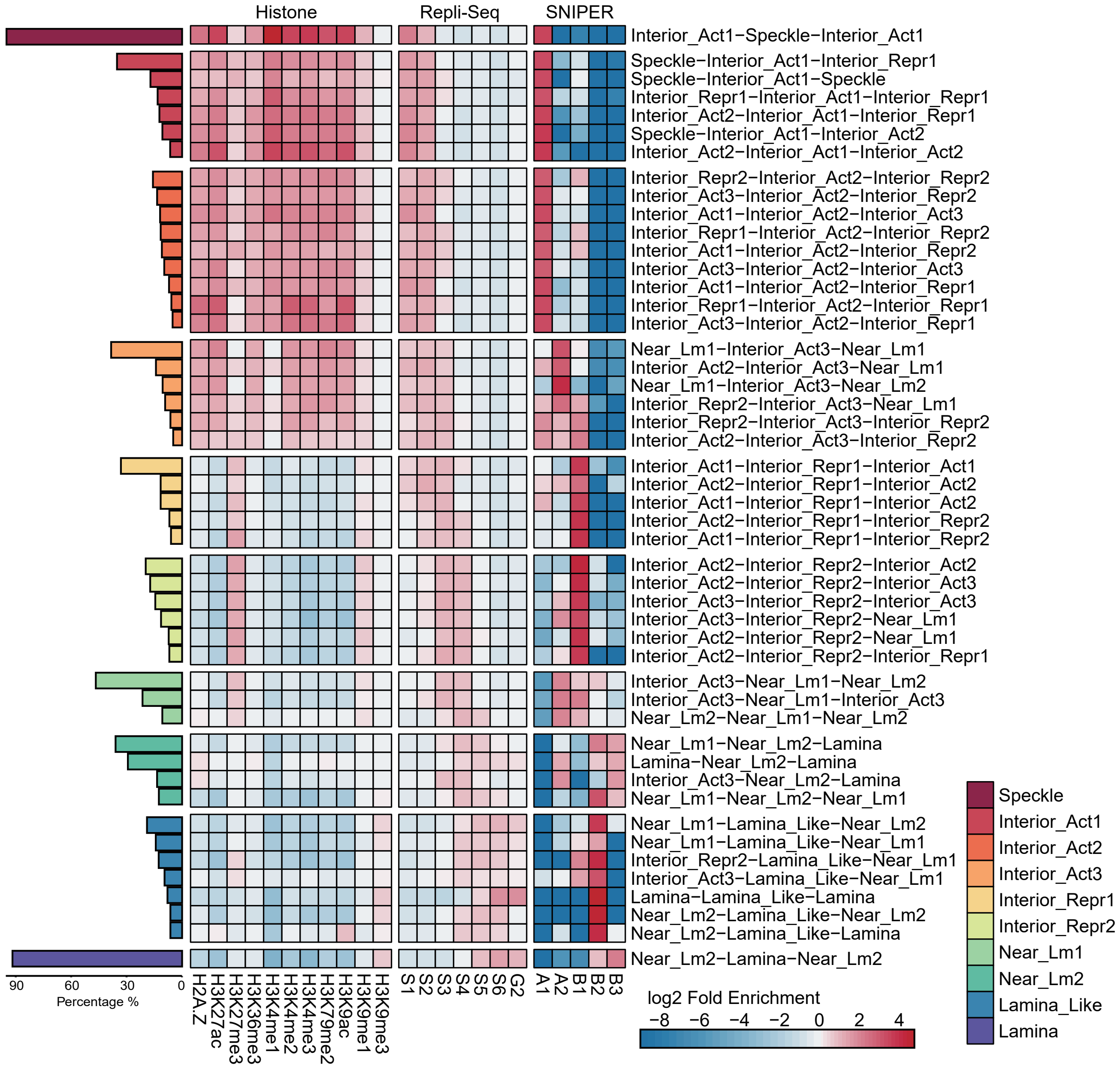


**Figure S12:** Heatmap showing the correlation between 3 consecutive SPIN states with histone mark signals, Repli-seq, and Hi-C subcompartments. The rows in the heat map represent different 3 SPIN patterns and are grouped by the SPIN states in the middle. Percentage of each 3 consecutive pattern for each SPIN state is shown on the left. Symmetrical 3 consecutive SPIN patterns are merged, and 3 consecutive SPIN patterns with length <100kb are not include. The heatmap shows the average fold enrichment of genomic signals on the middle SPIN state of each 3 consecutive pattern.

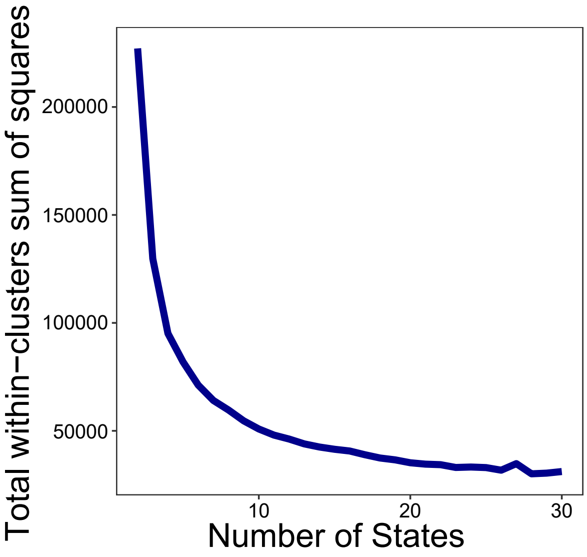


**Figure S13:** Determining the number of states. To determine optimal number of states, we first use $K$-means clustering algorithm to cluster 25kb bins based on TSA-seq and DamID signals. We choose different number of states (k), ranging from 2 to 30. Then we calculate the total within-clusters sum of squares for each cluster results.

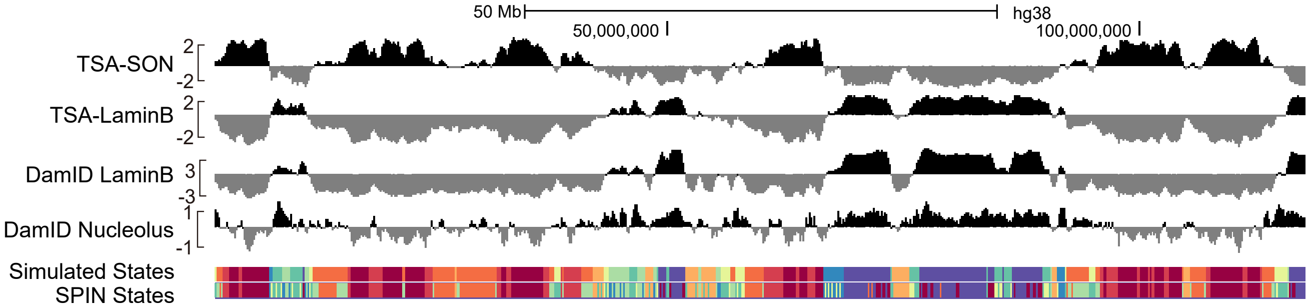


**Figure S14:** An example of the simulated data. Tracks shown here are (from top to bottom) the simulated TSA-seq SON, TSA-seq LaminB, DamID LaminB, DamID Nucleolus signal, the simulated states, and the SPIN states.

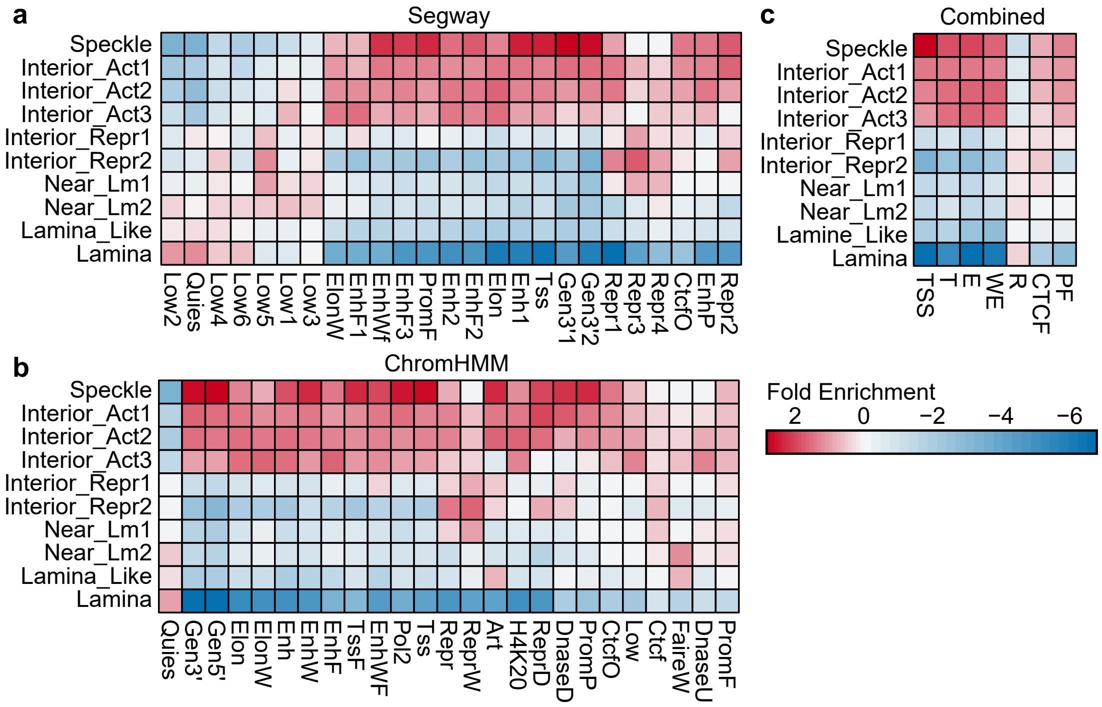


**Figure S15:** Comparison with ChromHMM and Segway chromatin states. The heatmap shows the correlation between SPIN states and the chromatin states from ChromHMM, Segway, and the combined segmentation. **a.** Comparison with Segway states in K562. **b.** Comparison with in K562. **c.** Comparison with combined states from Segway and ChromHMM. For each ChromHMM/Segway/Combined state, fold enrichment score on each SPIN state is calculated.

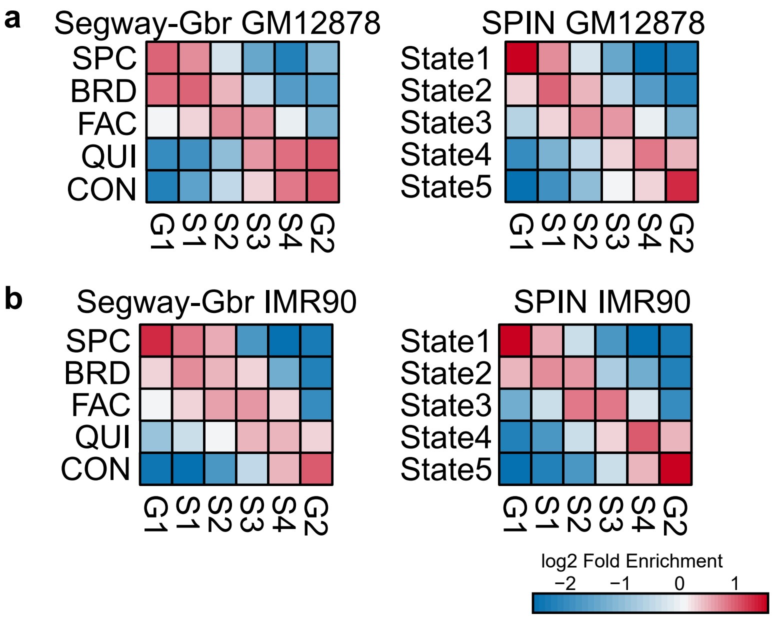


**Figure S16:** Comparison between SPIN and Segway-GBR. We used the same input data by Segway-GBR in GM12878 and IMR90 to run SPIN and set the number of states to 5. The heatmaps show the fold enrichment of Repli-seq signals in GM12878 and IMR90 stratified by the SPIN and Segway-GBR states, respectively. **a.** Comparison in GM12878. **b.** Comparison in IMR90.

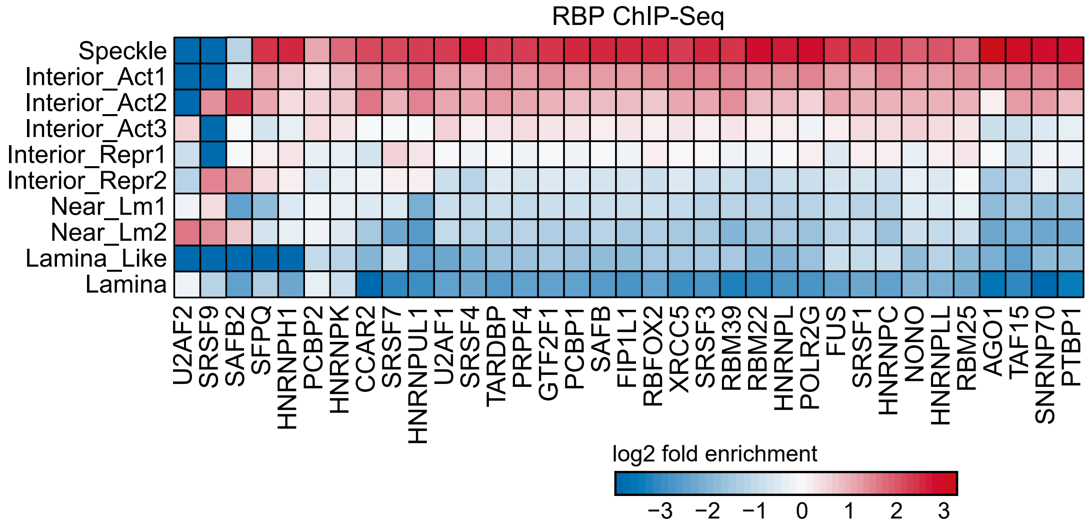


**Figure S17:** Heatmap showing RBP binding stratified by different SPIN states. The RBP ChIP-seq datasets were collected from (Xiao et al., 2019). log2 fold enrichment ChIP-seq peak density over expected is calculated.

| Data | Source | Identifier |
|---|---|---|
| TSA-seq SON | Chen et al. (2018) | GEO:GSE66019 |
| TSA-seq LaminB | Chen et al. (2018) | GEO:GSE66019 |
| DamID LaminB | Leemans et al. (2019) | 4DNEXZKHQQXY |
| DamID Nucleolus | This paper | - |
| Multi-fraction Repli-seq | This paper | - |
| *in situ* Hi-C | Rao et al. (2014) | GEO:GSE63525 |
| H2A.Z ChIP-Seq | ENCODE | ENCFF825TEU |
| H3K27ac ChIP-Seq | ENCODE | ENCFF094XCU |
| H3K27me3 ChIP-Seq | ENCODE | ENCFF366NNJ |
| H3K36me3 ChIP-Seq | ENCODE | ENCFF504HFV |
| H3K4me1 ChIP-Seq | ENCODE | ENCFF783QIW |
| H3K4me2 ChIP-Seq | ENCODE | ENCFF523HCI |
| H3K4me3 ChIP-Seq | ENCODE | ENCFF036HPL |
| H3K79me2 ChIP-Seq | ENCODE | ENCFF477LXF |
| H3K9ac ChIP-Seq | ENCODE | ENCFF390RCI |
| H3K9me1 ChIP-Seq | ENCODE | ENCFF823LIL |
| H3K9me3 ChIP-Seq | ENCODE | ENCFF559MMQ |
| K562 GRO-Seq | Niskanen et al. (2015) | GEO:GSE66448 |
| K562 LADs | 4DN data portal | 4DNESTAJJM3X |
| HCT116 LADs | 4DN data portal | 4DNES24XA7U8 |
| RPE-hTERT LADs | 4DN data portal | 4DNESHGTQ73M |
| HAP-1 LADs | 4DN data portal | 4DNESUK5H9Y8 |
| HFFc6 LADs | 4DN data portal | 4DNESXZ4FW4T |
| H1-hESC LADs | 4DN data portal | 4DNESXKBPZKQ |
| SNIPER Hi-C subcompartments | Xiong and Ma (2019) | - |
| Pol2 ChIA-PET | ENCODE | ENCSR000BZY |
| CTCF ChIA-PET | ENCODE | ENCSR000CAC |

**Table S1:** Datasets used in this paper.

| Region 1 | Region 2 | SV Type |
|---|---|---|
| chr1:6392018-6392019 | chr1:6558778-6558779 | germline_both_alleles dup |
| chr1:72300639-72300640 | chr1:72346157-72346158 | somatic del_l |
| chr1:172184314-172184315 | chr1:172646530-172646531 | somatic_post_aneuploid dup_l |
| chr1:174124133-174124134 | chr1:174195314-174195315 | somatic_post_aneuploid dup |
| chr1:209453861-209453862 | chr1:212914491-212914492 | somatic_post_aneuploid del_l |
| chr1:210154737-210154738 | chr1:210649177-210649178 | somatic_post_aneuploid del_l |
| chr10:37502218-37502219 | chr10:37578009-37578010 | somatic del_l |
| chr10:50358857-50358858 | chr10:50524669-50524670 | somatic_post_aneuploid del_l |
| chr10:86089577-86089578 | chr3:48186687-48186688 | somatic_pre_aneuploid/somatic_post_aneuploid |
| chr10:101843386-101843387 | chr10:102036378-102036379 | somatic_pre_aneuploid del_l |
| chr10:124952896-124952897 | chr10:125042150-125042151 | somatic_pre_aneuploid del_l |
| chr11:5312151-5312152 | chr11:5443878-5443879 | somatic_post_aneuploid del_l |
| chr11:54778070-54778071 | chr11:54756317-54756318 | somatic/somatic_post_aneuploid del_l |
| chr11:95025436-95025437 | chr11:95049343-95049344 | somatic dup |
| chr12:8208855-8208856 | chr12:8238297-8238298 | somatic_pre_aneuploid dup |
| chr13:80513898-80513899 | chr13:80895802-80895803 | somatic_post_aneuploid/somatic |
| chr13:80535269-80535270 | chr13:89786446-89786447 | somatic_post_aneuploid/somatic_pre_aneuploid del_l |
| chr13:80895894-80895895 | chr13:80896177-80896178 | somatic_post_aneuploid |
| chr13:91821754-91821755 | chr13:91823505-91823506 | somatic_post_aneuploid |
| chr13:93371154-93371155 | chr13:107848625-107848626 | somatic_post_aneuploid/somatic_pre_aneuploid del_l |
| chr13:107925598-107925599 | chr8:88686923-88686924 | somatic_post_aneuploid/somatic |
| chr13:108009064-108009065 | chr9:131280134-131280135 | somatic_pre_aneuploid/somatic_post_aneuploid |
| chr14:33997752-33997753 | chr14:34060109-34060110 | somatic_post_aneuploid dup |
| chr14:40381769-40381770 | chr2:23739093-23739094 | somatic_post_aneuploid/ |
| chr14:66304880-66304881 | chr14:66445743-66445744 | somatic_post_aneuploid del_l |
| chr14:66949670-66949671 | chr14:67186581-67186582 | somatic_post_aneuploid dup_l |
| chr14:96558592-96558593 | chr14:97172427-97172428 | somatic_post_aneuploid dup_l |
| chr15:76415871-76415872 | chr15:76673168-76673169 | somatic_post_aneuploid/germline_both_alleles del_l |
| chr16:20534460-20534461 | chr16:20597044-20597045 | somatic_pre_aneuploid dup |
| chr16:46689475-46689476 | chr16:46751233-46751234 | somatic_post_aneuploid/somatic |
| chr16:46692881-46692882 | chr16:46817414-46817415 | somatic |
| chr16:46978175-46978176 | chr16:47141183-47141184 | somatic dup |
| chr16:61021542-61021543 | chr16:61034696-61034697 | somatic_post_aneuploid |
| chr16:78226112-78226113 | chr6:38139520-38139521 | somatic_post_aneuploid |
| chr17:28573111-28573112 | chr17:28945867-28945868 | germline_both_alleles/somatic_post_aneuploid dup_l |
| chr17:46087894-46087895 | chr17:46356488-46356489 | somatic_post_aneuploid dup_l |
| chr17:66354134-66354135 | chr17:66503098-66503099 | somatic_post_aneuploid del_l |
| chr18:478328-478329 | chr18:3473273-3473274 | somatic_post_aneuploid |
| chr18:481781-481782 | chr18:21920738-21920739 | somatic_post_aneuploid/somatic del_l |
| chr18:3478988-3478989 | chr18:10860079-10860080 | somatic_post_aneuploid |
| chr18:7315202-7315203 | chr18:25949502-25949503 | somatic_post_aneuploid |
| chr18:8110158-8110159 | chr18:26776312-26776313 | somatic_post_aneuploid/ |
| chr18:8110681-8110682 | chr18:23730828-23730829 | somatic_post_aneuploid dup_l |
| chr18:41981480-41981481 | chr18:42007127-42007128 | somatic_post_aneuploid |
| chr19:14852281-14852282 | chr19:14903453-14903454 | somatic_post_aneuploid del_l |
| chr19:20413002-20413003 | chr19:20535164-20535165 | somatic_pre_aneuploid del_l |

**Table S2:** List of structural variations in K562 used in this work (combination of Li et al. (2016) and Dixon et al. (2018)).

| Region 1 | Region 2 | SV Type |
|---|---|---|
| chr2:112949812-112949813 | chr2:113117738-113117739 | somatic_post_aneuploid dup |
| chr2:144869687-144869688 | chr2:144979572-144979573 | somatic_post_aneuploid dup |
| chr2:148583522-148583523 | chr2:148585252-148585253 | somatic_post_aneuploid |
| chr22:20270018-20270019 | chr22:20271721-20271722 | somatic_post_aneuploid |
| chr22:20699515-20699516 | chr22:21716634-21716635 | somatic_post_aneuploid/somatic |
| chr22:22223455-22223456 | chr22:22333995-22333996 | somatic_post_aneuploid del_l |
| chr22:22585820-22585821 | chr4:4263840-4263841 | somatic_post_aneuploid |
| chr22:23290549-23290550 | chr9:130731760-130731761 | somatic_post_aneuploid/somatic_pre_aneuploid |
| chr22:23931954-23931955 | chr22:23969109-23969110 | somatic_post_aneuploid del_l |
| chr3:59757602-59757603 | chr3:60090194-60090195 | somatic_post_aneuploid del_l |
| chr3:60249783-60249784 | chr3:60526865-60526866 | somatic_post_aneuploid dup_l |
| chr3:60497197-60497198 | chr3:60556775-60556776 | somatic_post_aneuploid/somatic_pre_aneuploid dup |
| chr3:87865479-87865480 | chr3:87913904-87913905 | somatic_post_aneuploid del_l |
| chr3:138565637-138565638 | chr3:138613781-138613782 | somatic_post_aneuploid |
| chr3:160646056-160646057 | chr3:160982925-160982926 | somatic_post_aneuploid dup_l |
| chr4:58573762-58573763 | chr4:58674491-58674492 | somatic dup |
| chr4:69762251-69762252 | chr5:37296155-37296156 | somatic_post_aneuploid |
| chr4:157035097-157035098 | chr4:157111248-157111249 | somatic dup |
| chr4:159570179-159570180 | chr4:162695540-162695541 | somatic_pre_aneuploid del_l |
| chr5:22405253-22405254 | chr5:22450550-22450551 | somatic_post_aneuploid dup |
| chr5:54165099-54165100 | chr5:54266066-54266067 | somatic del_l |
| chr5:147086587-147086588 | chr5:147146778-147146779 | somatic del_l |
| chr5:163794104-163794105 | chr5:164558896-164558897 | somatic_post_aneuploid dup_l |
| chr6:16774234-16774235 | chr6:51870916-51870917 | somatic_post_aneuploid |
| chr6:29798048-29798049 | chr6:29953038-29953039 | germline_both_alleles/somatic_post_aneuploid dup |
| chr6:31651287-31651288 | chr6:31865935-31865936 | somatic_post_aneuploid dup_l |
| chr6:55961160-55961161 | chr6:55981915-55981916 | somatic del_l |
| chr6:68790254-68790255 | chr6:68825835-68825836 | somatic dup |
| chr6:71137307-71137308 | chr6:71169206-71169207 | somatic del_l |
| chr6:103289590-103289591 | chr6:103315010-103315011 | somatic del_l |
| chr6:160986407-160986408 | chr6:161246610-161246611 | somatic_post_aneuploid dup_l |
| chr7:8786866-8786867 | chr7:8826622-8826623 | somatic_pre_aneuploid del_l |
| chr7:36282546-36282547 | chr7:36701470-36701471 | somatic_post_aneuploid dup_l |
| chr7:79671139-79671140 | chr7:80139200-80139201 | somatic_post_aneuploid dup_l |
| chr7:98237080-98237081 | chr7:98284016-98284017 | somatic_pre_aneuploid del_l |
| chr7:111476851-111476852 | chr7:111773292-111773293 | somatic_post_aneuploid dup_l |
| chr8:2190308-2190309 | chr8:2212702-2212703 | somatic del_l |
| chr8:39374555-39374556 | chr8:39529710-39529711 | somatic_pre_aneuploid del_l |
| chr9:26589500-26589501 | chr9:38430029-38430030 | somatic/somatic_post_aneuploid dup_l |
| chr9:94542687-94542688 | chr9:97217791-97217792 | somatic_post_aneuploid del_l |
| chr9:112395416-112395417 | chr9:112652428-112652429 | somatic_post_aneuploid dup_l |
| chr9:120790476-120790477 | chr9:128341077-128341078 | somatic_post_aneuploid dup_l |
| chr9:120805644-120805645 | chr9:120872538-120872539 | somatic_post_aneuploid |
| chr9:120888023-120888024 | chr9:120908854-120908855 | somatic_post_aneuploid |
| chr9:128145977-128145978 | chr9:128147254-128147255 | somatic_post_aneuploid |
| chr9:128347614-128347615 | chr9:128702094-128702095 | somatic_post_aneuploid |

**Table S3:** (Continued) List of structural variations in K562 used in this work (combination of Li et al. (2016) and Dixon et al. (2018)).

| Accuracy | $K$-means | Gaussian HMM | SPIN |
|---|---|---|---|
| Speckle | 81% | 91% | 96% |
| Interior_Act | 77% | 85% | 91% |
| Interior_Repr | 54% | 64% | 73% |
| Near_Lm | 56% | 65% | 78% |
| Lamina | 71% | 80% | 86% |
| **Rand index** | 0.67 | 0.78 | 0.86 |

**Table S4:** Comparison of the genome segmentation results from $K$-means, Gaussian HMM, and SPIN on the simulated data. The prediction accuracy and rand index of each method are shown. Note that SPIN performed significantly better in all states.

| TF Cluster | GO Terms | Type | Gene Count | $p$-value |
|---|---|---|---|---|
| Cluster1 | GAF domain binding | MF | 11 | 9.92E-04 |
| | cellular response to L-ascorbic acid | BP | 74 | 1.26E-15 |
| | negative regulation of glutamine transport | BP | 74 | 1.26E-15 |
| | negative regulation of calcium-dependent cell-cell adhesion | BP | 36 | 6.01E-08 |
| Cluster2 | potassium channel inhibitor activity | MF | 81 | 1.13E-13 |
| | nucleotide diphosphatase activity | MF | 68 | 1.11E-12 |
| | GPI-linked ephrin receptor activity | MF | 55 | 1.09E-11 |
| | PH domain binding | MF | 64 | 4.19E-10 |
| | serotonin binding | MF | 76 | 4.43E-10 |
| | olfactory bulb interneuron development | BP | 92 | 3.37E-14 |
| | establishment of integrated proviral latency | BP | 79 | 4.78E-14 |
| | establishment of viral latency | BP | 80 | 6.60E-14 |
| | radial glia guided migration of Purkinje cell | BP | 73 | 7.05E-14 |
| | viral latency | BP | 80 | 1.06E-13 |
| | nonhomologous end joining complex | CC | 72 | 2.82E-11 |
| | growth cone membrane | CC | 80 | 2.03E-10 |
| | DNA ligase IV complex | CC | 44 | 3.72E-10 |
| | insulin receptor complex | CC | 54 | 1.67E-08 |
| | DNA-dependent protein kinase-DNA ligase 4 complex | CC | 50 | 1.42E-07 |
| Cluster3 | taurine:sodium symporter activity | MF | 34 | 1.24E-11 |
| | methionine adenosyltransferase activity | MF | 31 | 1.13E-10 |
| | sarcosine dehydrogenase activity | MF | 34 | 2.32E-10 |
| | recombinase activity | MF | 28 | 1.04E-09 |
| | glycerate dehydrogenase activity | MF | 28 | 1.04E-09 |
| | heparan sulfate proteoglycan catabolic process | BP | 60 | 2.98E-17 |
| | regulation of proton-transporting ATPase activity | BP | 41 | 1.57E-12 |
| | taurine transport | BP | 34 | 1.24E-11 |
| | epidermal cell fate specification | BP | 28 | 1.04E-09 |
| | arterial endothelial cell fate commitment | BP | 28 | 1.04E-09 |
| | centrosomal corona | CC | 31 | 1.95E-09 |

**Table S5:** GO analysis of genes near ChIP-seq peaks in different TF clusters based on the SPIN states. ChIP-seq peaks on each clusters are merged and used as input for GREAT to calculate GO term enrichment.
